# Effects of dopamine receptor antagonism and amphetamine-induced psychomotor sensitization on sign- and goal-tracking after extended training

**DOI:** 10.1101/848150

**Authors:** Shaun Yon-Seng Khoo, Alexandra Uhrig, Anne-Noël Samaha, Nadia Chaudhri

## Abstract

The dopamine system is important for incentive salience attribution, where motivational value is assigned to conditioned cues that predict appetitive reinforcers. However, the role of dopamine in this process may change with extended training. We tested the effects of dopamine D1-like and D2-like receptor antagonism on the expression of sign-tracking and goal-tracking conditioned responses following extended Pavlovian conditioned approach (PCA) training. We also tested if amphetamine-induced psychomotor sensitization accelerates the enhanced acquisition of sign-tracking that is observed with extended training. In experiment 1, 24 male Long-Evans rats received 20 PCA sessions in which one lever (CS+, 10 s) predicted 0.2 mL sucrose (10%, w/v) delivery and the other lever (CS−) did not. SCH-23390 (D1-like antagonist) or eticlopride (D2-like antagonist) were administered before non-reinforced behavioural tests at doses of 0, 0.01, and 0.1 mg/kg (s.c.). In experiment 2, rats received vehicle or 2 mg/kg amphetamine (i.p.) for 7 days (n = 12/group). Ten days later, they received 16 PCA training sessions. Both doses of SCH-23390 reduced sign- and goal-tracking, but also reduced locomotor behaviour. A low dose of eticlopride (0.01 mg/kg) selectively reduced goal-tracking, without affecting sign-tracking or locomotor behaviour. Amphetamine produced psychomotor sensitization, and this did not affect the acquisition of sign- or goal-tracking. Following extended PCA training, dopamine D2-like receptor activity is required for the expression of goal-tracking but not sign-tracking. Psychomotor sensitization to amphetamine did not impact incentive salience attribution; however, more selective manipulations of the dopamine system may be needed.

## Introduction

Environmental cues that predict appetitive reinforcers can acquire incentive salience, whereby motivational value is assigned to the cue (Berridge, 2007; Chow et al., 2016; Flagel et al., 2011, 2007; Fraser & Janak, 2017). Incentive salience has been studied using a Pavlovian conditioned approach (PCA) or ‘autoshaping’ task, where the brief insertion and then retraction of a lever conditioned stimulus (CS) is paired with an appetitive, unconditioned stimulus (US). When the CS is presented, sign-tracking animals preferentially approach and interact with the CS, while goal-tracking animals preferentially approach the location where the US will be presented (Berridge & Robinson, 2003; Flagel et al., 2007). Both sign- and goal-trackers learn the CS-US association, but only sign-trackers assign motivational value to the CS.

Most studies characterise animals’ sign- and goal-tracking behaviour after a short 5 day period of PCA training (Flagel et al., 2007; Meyer et al., 2012; Robinson & Berridge, 2008), however there is evidence that longer periods of PCA training result in behavioural and neuropharmacological differences. We have reported that extended PCA training can produce a shift from goal-tracking to sign-tracking. In one study, goal-tracking responses to an alcohol-associated CS in rats peaked between sessions 7 and 10 of PCA training before decreasing, while the number of sign-tracking responses continued to increase over 27 PCA sessions (Srey et al., 2015). In a subsequent, large-scale analysis of data from 5 different experiments, this shift from goal-to sign-tracking behaviour emerged as a significant behavioural phenotype (Villaruel & Chaudhri, 2016). However, it is currently unclear which neural systems could play a role in mediating the shift from goal-to sign-tracking behaviour that emerges after extended PCA training.

Dopamine signalling has been heavily implicated in the acquisition and expression of incentive salience in studies using the PCA task (see Table 1) and it may be altered by extended training. Systemic injections of a non-selective dopamine antagonist during PCA training impaired the expression of both sign-tracking and goal-tracking, and also blocked the acquisition of sign-tracking but not goal-tracking after 7 PCA sessions (Flagel et al., 2011). Moreover, several studies suggest that dopamine signalling at D1-like receptors (D1 and D5; hereafter, D1) in the nucleus accumbens core is particularly important in the attribution of incentive salience to an appetitive CS after 3-8 PCA sessions (Chow et al., 2016; Clark et al., 2013; Dalley et al., 2005; Flagel et al., 2011; Roughley & Killcross, 2019; Saunders & Robinson, 2012). However, it has also been shown that extended PCA training can reduce the extent to which sign-tracking relies on dopamine signalling. Sign-trackers show greater expression of D1 receptor mRNA during the first session of PCA training than goal-trackers, while goal-trackers exhibit greater tyrosine hydroxylase, dopamine transporter and D2 receptor mRNA expression than sign-trackers in later sessions (Flagel et al., 2007). Moreover, measurement of CS-elicited dopamine release over the course of extended training has revealed that, in sign-trackers, dopamine release peaks by the 4th training session and then diminishes by the 15th PCA session (Clark et al., 2013). This change in dopamine signalling over time may explain why administering eticlopride during a 14-day PCA protocol impaired goal-tracking (Chow et al., 2016), but administering eticlopride after 4 PCA sessions impaired sign-tracking (Lopez et al., 2015). These results suggest that extended training may alter the role of the dopamine system in sign-tracking and goal-tracking, but the relative contributions of D1-like and D2-like receptors after extended training are unclear.

**Table 1.** Summary of previous studies on the role of dopamine in acquisition and expression of sign- and goal-tracking

| Animals (Housing)* | Design | Manipulation | Result | Reference |
| --- | --- | --- | --- | --- |
| Male Sprague-Dawley rats (Individual housing) | CS+, 8 sessions, 25 trials/session, banana pellet reinforcer | D1&D2: Acb core flupenthixol before test | Impaired expression of sign-tracking | Saunders & Robinson (2012) |
| Male Sprague-Dawley rats (Pair-housed) | CS+, 5 sessions, 25 trials/session, food pellet reinforcer | D1&D2: *In situ* hybridization after session 1 or 5 | Session 1: D1 mRNA greater in sign-trackers than goal-trackers<br>Session 5: Tyrosine hydroxylase, dopamine transporter, D2 mRNA greater in goal-trackers than sign-trackers | Flagel et al. (2007) |
| Male Sprague-Dawley rats bred to respond to novelty (Individual housing) | CS+, 7 sessions, 25 trials/session, food reinforcer | D1&D2: Flupenthixol before training<br>DA: Fast scan cyclic voltammetry | Flupenthixol prevented expression of sign and goal-tracking. Only acquisition of sign-tracking was blocked. | Flagel et al. (2011) |
| Male Sprague-Dawley rats (Individual housing) | CS+, 15 sessions, 25 trials/session, food pellet reinforcer | DA: Fast scan cyclic voltammetry in sign-trackers | Peak CS-evoked dopamine rises early in training, then diminishes | Clark et al. (2013) |
| Male Lister rats (Pair-housed) | CS+/CS-, 3 sessions, 50 trials/session, food pellet reinforcer | D1: Acb Core SCH-23390 after each session | Impaired acquisition | Dalley et al. (2005) |
| Male Sprague-Dawley rats (Individual housing) | Lever-CS+/Tone-CS+, 14 sessions, 16 trials per CS/session, food pellet reinforcer | D1: SCH-23390, i.p. intermittent pretreatment during training | Impaired acquisition of sign-tracking | Chow et al. (2016) |
| Male Wistar rats (Group-housed) | CS+, 7 sessions, 28 trials/session, food pellet reinforcer | D1: SCH-23390, i.p. pretreatment during training | Impaired acquisition of sign- and goal-tracking | Roughley & Killcross (2019) |
| Male Sprague-Dawley rats (Individual housing) | CS+, 5 or 15 sessions, 25 trials/session, food pellet reinforcer | D1: SCH-23390, i.p. before test | Impaired expression of sign-tracking | Clark et al. (2013) |
| Male Sprague-Dawley rats (Individual housing) | CS+/CS-, 9 sessions, 15 trials/session, sucrose pellet reinforcer | D2: Haloperidol or olanzapine, i.p. pretreatment during training | Impaired acquisition of sign-tracking, but not goal-tracking | Danna & Elmer (2010) |
| Male Sprague-Dawley rats (Pair-housed) | CS+, 5 sessions, 25 trials/session, banana-flavoured food pellet reinforcer | D2: Haloperidol via minipump or via daily s.c. injections | Acute amphetamine-induced increase in goal-tracking was prevented after haloperidol (minipump or s.c.) discontinuation | Bédard et al. (2011) |
| Male Wistar rats (Group-housed) | CS+, 7 sessions, 28 trials/session, food pellet reinforcer | D2: Eticlopride, i.p. pretreatment during training | Impaired acquisition of sign-tracking and expression of goal-tracking | Roughley & Killcross (2019) |
| Male Long-Evans rats (Group-housed) | CS+, 4 sessions, 25 trials/session, food pellet reinforcer | D2: Eticlopride, i.p. before test sessions | Impaired expression of sign- and goal-tracking | Lopez et al. (2015) |
| Male Sprague-Dawley rats (Individual housing) | Lever-CS+/Tone-CS+, 14 sessions, 16 trials per CS/session, food pellet reinforcer | D2: Eticlopride, i.p. intermittent pretreatment during training | Impaired expression of sign- and goal-tracking | Chow et al. (2016) |
| Male Sprague-Dawley rats (Pair/Group-housed) | CS+, 7 sessions, 25 trials/session, banana pellet reinforcer | D2: 7OH-DPAT, pramipexole, or raclopride i.p. before test sessions | Both agonists (7OH-DPAT, pramipexole) and antagonist (raclopride) impaired expression of sign- and goal-tracking | Fraser et al. (2016) |

\* Individually housed: 1 per cage. Pair-housed: 2 per cage. Group-housed: Typically, 3-4 per cage.

The aim of our first study was therefore to compare the effects of dopamine D1-like and D2-like receptors in the expression of sign- and goal-tracking behaviour in rats that had received extended (20 days) PCA training, as these effects have not been examined directly in other studies. We gave rats extended PCA training and classified them as sign-trackers, goal-trackers or intermediates using criteria developed by Meyer et al. (2012). We then tested the dose-dependent effects of the selective D1-like and D2like receptor antagonists, SCH-23390 and eticlopride, on the expression of sign- and goal-tracking behaviour to verify whether dopamine receptor signalling was necessary for the expression of incentive salience after extended PCA training. As an additional behavioural control, we examined the effects of these antagonists on locomotor activity tested in an open field arena.

Extended training can also produce a shift in phenotypes and several studies have attempted to examine the effects of drug-induced psychomotor sensitization on changes to PCA phenotypes over time, with conflicting results. Psychomotor sensitization develops with repeated exposure to drugs, such as amphetamine, and it renders animals more sensitive to drug-induced increases in psychomotor activity, incentive motivation, and dopamine neurotransmission (Downs & Eddy, 1932; Kalivas & Stewart, 1991; Robinson & Becker, 1986; Segal & Mandell, 1974). Sensitization can enhance cue-induced dopamine release, thereby imbuing cues with increased incentive salience (Bradberry, 2007; Leyton, 2007; Leyton & Vezina, 2013; Robinson & Berridge, 1993). If sensitization increases the incentive salience of cues, then animals might be expected to approach reward cues, or sign-track, more. However, prior studies have examined the influence of amphetamine-induced psychomotor sensitization on sign- and goal-tracking, with conflicting results. Some found enhanced sign-tracking responses (Robinson et al., 2015; Wyvell & Berridge, 2001) while others found enhanced goal-tracking responses (Simon et al., 2008). These studies may suggest opposing effects on incentive salience, but are difficult to directly compare due to procedural differences. For example, protocols differed in whether sensitization was induced before or after Pavlovian conditioning and rats variously received 7, 8 or 14 PCA sessions in different studies (Robinson et al., 2015; Simon et al., 2008; Wyvell & Berridge, 2001). Despite the difficulty in interpreting these conflicting results, these studies suggest that psychomotor sensitization to amphetamine has the potential to influence changes in sign-tracking and goal-tracking over time. Moreover, it is currently unknown whether amphetamine-induced psychomotor sensitization, prior to PCA training, might alter the rate at which animals acquire a sign-tracking phenotype over time (Srey et al., 2015; Villaruel & Chaudhri, 2016). To address this, our second objective was to determine if amphetamine-induced psychomotor sensitization would accelerate the acquisition of sign-tracking which can occur with extended training

In Experiment 2, we therefore examined the effect of amphetamine sensitization on the acquisition of sign-tracking over time. Prior to PCA training, rats received a series of vehicle or amphetamine injections. We hypothesised that, over the course of 16 PCA sessions, rats that had previously received amphetamine injections would acquire sign-tracking more rapidly than control rats do.

## Materials and Methods

### Animals

Subjects were 48 experimentally naïve male Long-Evans rats weighing 220-240 g on arrival, corresponding to approximately 6-10 weeks of age (Charles River). Only male rats were used because procedures were not yet optimised in female rats in our laboratory. Rats were initially pair-housed in plastic cages (44.5 × 25.8 × 21.7 cm) containing Teklad Sani Chip bedding, a nylon bone chew toy, a plastic tunnel, and shredded paper in a climate-controlled (21°C) vivarium on a 12 h:12 h light:dark cycle (lights on at 7am). Rats acclimated to the colony room over at least 3 days before being singly-housed and handled for 7 days. Single housing was to enable accurate measurement of home-cage sucrose consumption and housing conditions, including enrichment, were otherwise unchanged. Rats remained singly housed for the rest of the experiment to keep housing conditions consistent throughout the study. Rats had free access to food and water in their home-cages throughout the experiments. All procedures were approved by the Animal Research Ethics Committee at Concordia University and accorded with guidelines from the Canadian Council on Animal Care. See Table 2 for details concerning rats, reagents, and equipment.

**Table 2.**
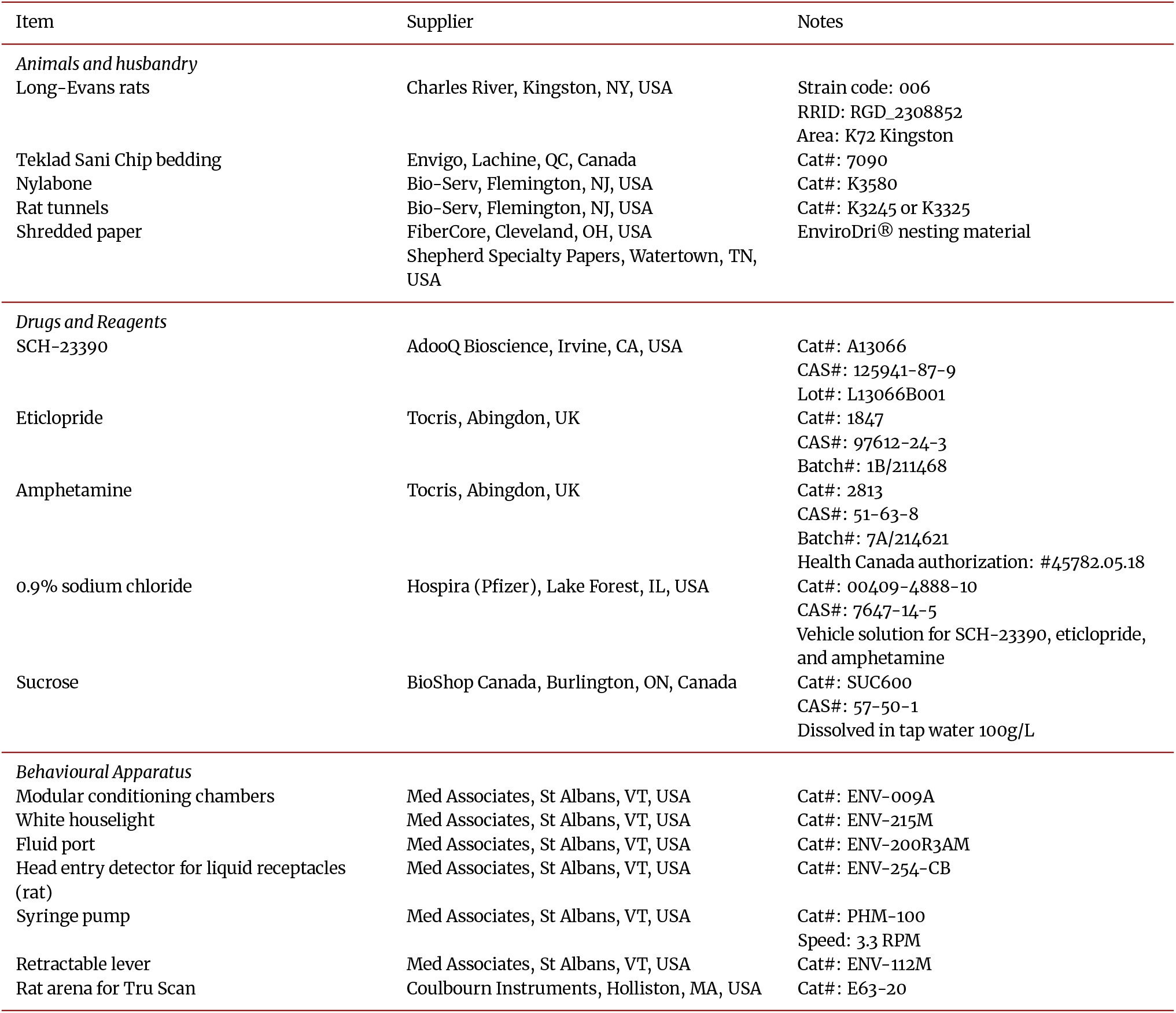
Materials and Supplier Details

### Apparatus

Behavioural training was conducted using 12 identical conditioning chambers (30.5 x 31.8 x 29.2 cm, Med Associates). Each chamber was contained within a sound-attenuating cubicle with a fan to provide ventilation and background noise (70-75 dB). Each chamber had a white houselight in the center near the ceiling of the left wall (as viewed by the experimenter). The right wall had a fluid port with infrared detectors located above the floor. A 20 mL syringe was placed on a syringe pump outside the cubicle and connected to the port with polyethylene tubing. Retractable levers on either side of the port served as conditioning stimuli. A PC running Med-PC IV controlled stimulus presentations and recorded responses. For locomotor behaviour, we used four 39 x 42 x 50 cm open field arenas (Coulbourn Instruments) housed in sound attenuating boxes and Tru Scan 2.0 software to compute locomotion.

### Experiment 1: Effect of dopamine receptor antagonism on the expression of sign- and goal-tracking

#### Home-cage sucrose

To familiarize rats (n = 24) with sucrose, they received 48 h of unrestricted access to 10% sucrose in the home-cage. A pre-weighed bottle containing 90 mL sucrose and a bottle of water were placed on the home-cage. After 24 h, bottles were re-weighed, refilled, and replaced for 24 h. Rats consumed all, or nearly all, the sucrose.

#### Habituation

Rats were then habituated to the conditioning chambers. On day 1, they were exposed to the testing room for 20 min where they were handled and weighed. On day 2, rats were placed in the conditioning chambers, where after a programmed 2-min delay the houselights were switched on for a 20 min session in which port entries were counted, but had no programmed consequences.

#### Pavlovian conditioned approach

Rats then underwent 20 PCA sessions. Each session began with a 2-min delay, after which the houselight was switched on to indicate session onset. Each session involved 20 trials, with 10 CS+ trials (paired with sucrose) and 10 CS− trials (no sucrose). Each trial consisted of a 10-s Pre-CS interval, a 10-s CS lever presentation, and a 10-s Post-CS interval. One of the two levers was designated as the CS+ lever, while the other was the CS− lever. Levers were counterbalanced so that for half of the rats the CS+ lever was on the left of the fluid port and for the remainder the CS+ lever was on the right of the fluid port. For a CS+ trial, but not a CS− trial, 6 s of syringe pump operation began at the onset of the Post-CS interval (i.e., immediately after retraction of the CS+ lever) to deliver 0.2 mL of sucrose into the fluid port for oral consumption. The inter-trial intervals (ITI), which did not include the Pre-CS, CS, or Post-CS intervals, were set at 60, 120, or 180 s (mean ITI duration = 120 s). The ITI durations and order of CS+ and CS− trials were randomized.

For each session, a PCA score was calculated from response bias, probability difference, and latency score, as defined in Table 3 (Meyer et al., 2012). Rats were classified as sign-trackers if their mean PCA score was ≥ 0.5 for PCA sessions 19 and 20. Rats with PCA scores ≤ −0.5 were classified as goal-trackers and rats with PCA scores between −0.5 and 0.5 were intermediates (Ahrens et al., 2016; Meyer et al., 2012).

**Table 3.** Definitions of Key Variables

| Variable | Definition |
| --- | --- |
| *Pavlovian Conditioned Approach* |  |
| PCA Score | (Response Bias + Latency Score + Probability Difference) ÷ 3 |
| Response Bias | (Lever Activations – Port Entries) ÷ (Lever Activations + Port Entries) |
| Probability Difference | (Trials with Lever Activations – Trials with Port Entries) ÷ Number of Trials |
| Latency Score | (Mean Port Entry Latency – Mean Lever Activation Latency) ÷ CS Duration |
| *Locomotor Behaviour* |  |
| Distance Travelled | Sum of the coordinate changes during the session (cm) in the floor plane. |
| Movement Episodes | The number of episodes of movement in the floor plane, as defined by continuous coordinate changes without resting for at least one second. |
| Center Time | The amount of time spent in the center of the arena, as defined by the animal's coordinates being at least 2.5 beam-widths (6.25 cm) away from the chamber walls. |
| Corner Time | Time spent within 4 beams (10 cm) of each corner. |
| Center Distance | Sum of the coordinate changes during the session (cm) that occurred at least 2.5 beam-widths (6.25 cm) away from the chamber walls. |
| Stereotypy Time | The total time spent engaging in movements that resulted in coordinate changes of less than 1 beam (2.5 cm) and were less than 2 s apart. Time spent engaging in such restricted movements is potentially suggestive of stereotypy. |

#### Dopamine antagonist tests

We tested the effect of dopamine receptor antagonists on the expression of conditioned responding in the absence of sucrose (extinction conditions), with dose order counterbalanced using a within-subjects, Latin square design.

First, to test the effect of the D1 antagonist SCH-23390, rats received an injection of saline vehicle, 0.01 mg/kg, or 0.1 mg/kg SCH-23390 (1 mL/kg, s.c.) 15 min before a test session. Test sessions were identical to training sessions, but no syringes were placed on the pump and no sucrose was delivered. Rats received at least one day of PCA training between tests to allow them to return to baseline levels of responding.

After testing SCH-23390, the same procedure was repeated, but rats received saline vehicle, 0.01 mg/kg, or 0.1 mg/kg eticlopride. Previous studies have shown 0.01 mg/kg is a behaviourally effective dose for SCH-23390 or eticlopride (Chow et al., 2016; Sciascia et al., 2014) and in a pilot test conducted in the same rats we found that a lower dose (0.001 mg/kg SCH-23390 or eticlopride) had no effect on sign- or goal-tracking.

#### Locomotor activity test in an open field arena

In the same rats, we examined the effects of SCH 23390 and eticlopride on locomotor behaviour in an open field arena (see Table 3 for definition of variables). To prevent confounds, such as reduced locomotor behaviour across repeated tests, rats were tested using a between-subjects design, where rats received a single dose of each compound, and all rats were tested on the same day (1 day/compound). On day 1, rats were exposed to the open field arena for a 45 min habituation session. The next day, rats were randomly allocated to receive vehicle, 0.01 mg/kg, or 0.1 mg/kg SCH-23390 (1 mL/kg, s.c.) 15 min before a test in which we measured locomotor activity in the open field arena (n = 8/dose). On a separate day, rats were randomly allocated to receive vehicle, 0.01 mg/kg, or 0.1 mg/kg eticlopride (1 mL/kg, s.c., n = 8/dose) 15 min before a locomotor activity test.

### Experiment 2: Effect of amphetamine-induced psychomotor sensitization on the acquisition of sign- and goal-tracking

#### Amphetamine exposure

We gave rats repeated amphetamine injections in order to induce psychomotor sensitization. Since amphetamine sensitization can be context-specific (Badiani et al., 1995a,b; Crombag et al., 2001, 2000), we exposed rats to both the open field arena as well as the conditioning chamber after each amphetamine injection, such that amphetamine’s effects were paired with both contexts. On day 1, all rats (n = 24) were habituated to the procedure with a saline injection (1 mL/kg, i.p.) immediately before placement into an open field arena for 30 min, followed by placement into a conditioning chamber for 20 min. When in the conditioning chamber, entries into the fluid port were recorded, but no stimuli were presented. Rats were then randomly allocated (n = 12 /group) to receive saline vehicle (1 mL/kg) or amphetamine (2 mg/kg, 1 mL/kg, i.p.) for the next 7 consecutive days, which were otherwise identical to day 1. Doses were based on previous studies (Robinson et al., 2015).

#### Incubation of amphetamine-induced psychomotor sensitization and home-cage sucrose exposure

For the first 7 days after amphetamine or vehicle exposure, rats were left undisturbed in their home cages except for normal husbandry activities. On days 8-10, rats received 48 h of home-cage sucrose exposure, as described in Experiment 1.

#### Pavlovian conditioned approach

Behavioural training began on the 11th day after the last amphetamine/vehicle injection. Rats received 14 PCA sessions in the conditioning chambers using procedures that were identical to those described for Experiment 1.

#### Psychomotor sensitization test

After PCA training, rats were tested for sensitization to the psychomotor activating effects of amphetamine. All rats received a 0.75 mg/kg, 1 mL/kg, i.p. amphetamine challenge (Robinson et al., 2015) immediately before a 30 min locomotor activity test in an open field arena.

### Statistical Analysis and Material Availability

Data were analysed using SPSS 24 (IBM, NY, USA). ANOVA with Bonferroni-corrected post-hoc comparisons and t-tests were used. Greenhouse-Geisser corrections were applied to degrees of freedom following a significant Mauchly’s test of sphericity with ε< 0.75. Following violations of ANOVA assumptions for some locomotor measures in Experiment 1, the Kruskal-Wallace test was used. Raw data and Med-PC code are available on Figshare (Khoo et al., 2021).

## Results

### Experiment 1: Effect of dopamine receptor antagonism on the expression of sign- and goal-tracking

Following the acquisition of PCA, we identified the behavioural phenotypes of all rats based on PCA scores averaged across sessions 19 and 20 (Villaruel & Chaudhri, 2016). Fourteen rats were identified as sign-trackers, 6 as goal-trackers and 4 as intermediates. Conditioned responses obtained across 20 PCA sessions in these rats are shown in Figure A.1, with statistical results in Table A.1.

#### SCH-23390

Systemic administration of a dopamine D1-like receptor antagonist following extended PCA training reduced the expression of sign-tracking in sign-trackers and intermediates, reduced the expression of goal-tracking in goal-trackers, and reduced ITI port entries in all phenotypes (Figures 1a-c).

**Figure 1.**
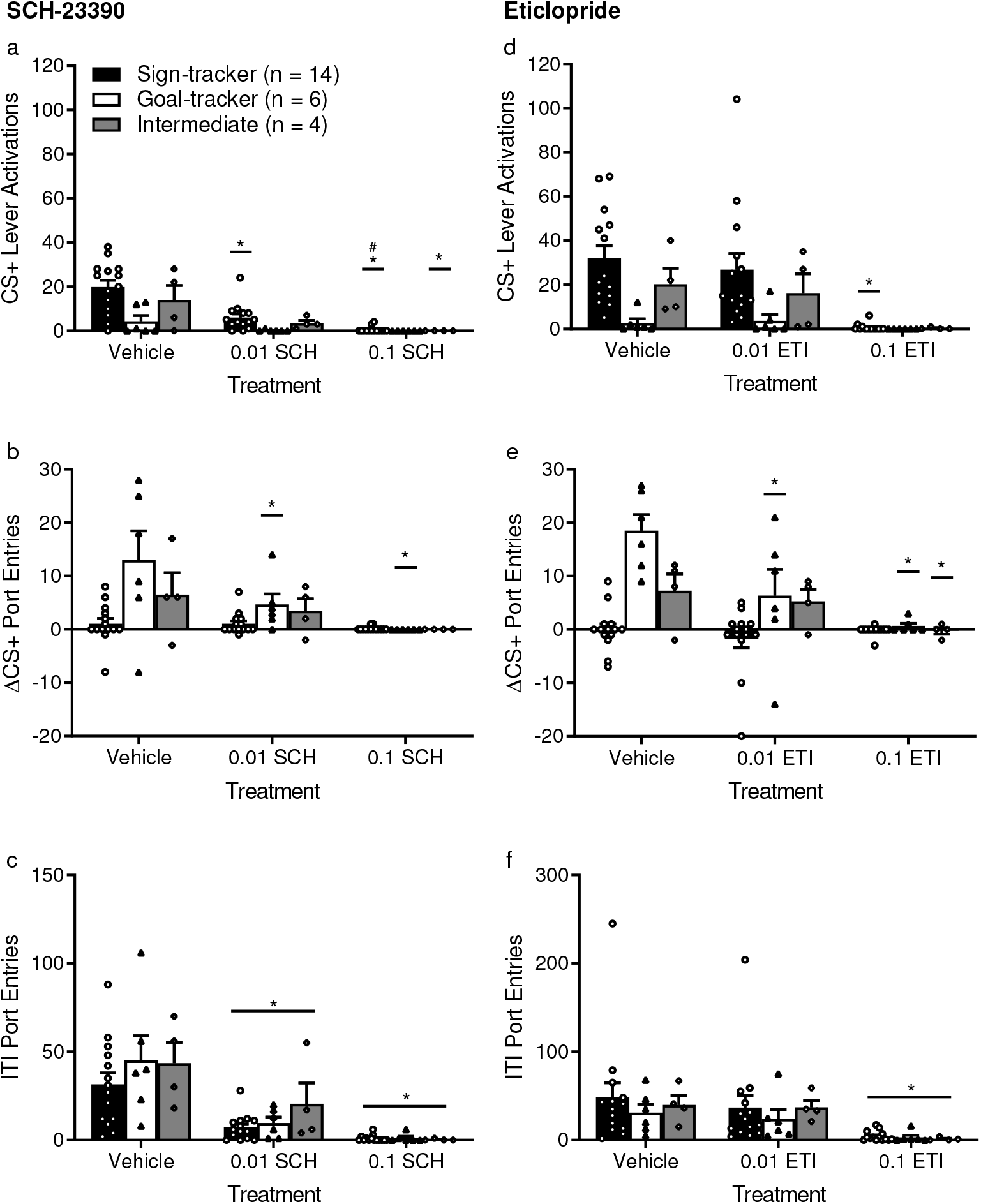
Antagonism of dopamine D1-like receptors reduced goal-tracking, sign-tracking and port entries during the intertrial interval (ITI), while dopamine D2-like receptor antagonism selectively reduced goal-tracking by goal-trackers. (a) The selective dopamine D1 receptor antagonist, SCH-23390, dose-dependently reduced CS+ lever activations by sign-trackers (n = 14). CS+ lever activations by intermediates (n = 4) were reduced at a dose of 0.1 mg/kg. (b) SCH-23390 dose-dependently reduced ΔCS+ port entries (CS+ port entries – Pre-CS+ port entries) in goal-trackers (n = 6). (c) SCH-23390 dose-dependently reduced ITI port entries across phenotypes. (d) The selective dopamine D2 receptor antagonist, eticlopride, reduced CS+ lever activations by sign-trackers at a dose of 0.1 mg/kg. (e) Eticlopride significantly reduced ΔCS+ port entries made by goal-trackers at both doses and reduced ΔCS+ port entries by intermediates at 0.1 mg/kg. (f) Eticlopride did not have significant effects on ITI port entries at a dose of 0.01 mg/kg but reduced ITI port entries across phenotypes at 0.1 mg/kg. ITIs comprised 42 min of each session. Testing was conducted under extinction conditions (no sucrose was delivered). Data are means ± SEM. * *p* < 0.05 for Bonferroni corrected post-hoc comparisons compared to vehicle. # *p* < 0.05 for Bonferroni corrected post-hoc comparison compared to 0.01 mg/kg.

CS+ lever activations (Figure 1a) was our behavioural index of sign-tracking. Because the levers were only extended as a CS, unadjusted lever activations were used rather than ΔCS lever activations (as will be described for port entries below). CS+ lever activations differed by phenotype [F(2,21) = 5.09, *p* = 0.016] and dose [F(1.347,28.294) = 20.441, *p* < 0.001, ε= 0.674] with a significant interaction [dose × phenotype, F(2.695,28.294) = 3.154, *p* = 0.045]. In sign-trackers, both 0.01 and 0.1 mg/kg of SCH-23390 reduced CS+ lever activations compared to vehicle (*p*’s < 0.001) and 0.1 mg/kg of SCH 23390 further reduced CS+ lever activations relative to 0.01 mg/kg SCH-23390 (*p* = 0.001). In intermediates, SCH-23390 reduced CS+ lever activations at the 0.1 mg/kg dose (*p* = 0.047), but not 0.01 mg/kg (*p* = 0.126), relative to vehicle. There were no differences between doses for goal-trackers (*p*’s ≥ 0.914).

ΔCS+ port entries (Figure 1b) was our behavioural index of goal-tracking. This measure considers differences in baseline levels of port entry behaviour by subtracting PreCS+ port entries from CS+ port entries (Chaudhri et al., 2013; Khoo et al., 2019; Panayi & Killcross, 2018). ΔCS+ port entries varied according to SCH-23390 dose [F(1.271,26.687) = 9.961, *p* = 0.002, ε= 0.635] and phe-notype [F(2,21) = 5.904, *p* = 0.009] and there was a significant dose × phenotype interaction [F(2.542,26.687) = 3.802, *p* = 0.027]. In goal-trackers, SCH-23390 decreased ΔCS+ port entries at the 0.01 mg/kg dose, compared to vehicle (*p* = 0.041), and the 0.1 mg/kg dose compared to vehicle (*p* = 0.002) and 0.01 mg/kg (*p* = 0.006). ΔCS+ port entries in sign-trackers and intermediates was not significantly affected by SCH-23390.

SCH-23390 dose-dependently reduced ITI port entries (Figure 1c]). There was a main effect of dose [F(1.233,25.899) = 28.072, *p* < 0.001, ε= 0.617], but no effect of phenotype [F(2,21) = 1.244, *p* = 0.309] or dose × phenotype interaction [F(2.467,25.899) = 0.686, *p* = 0.542]. Both SCH-23390 doses reduced ITI port entries (0.01 mg/kg vs. vehicle, *p* = 0.001; 0.1 mg/kg vs vehicle and 0.01 mg/kg, *p* < 0.001). Effects on the CS− and Post-CS periods are shown in Figure A.2 and effects on probability of and latency to CS+ lever activations and port entries and PCA score are presented in Figure A.3, with statistical results in Tables A.2 and A.3, respectively.

#### Eticlopride

Figures 1d-f show effects of the dopamine D2-like receptor antagonist, eticlopride. Systemic administration of low dose (0.01 mg/kg) eticlopride following extended PCA training reduced the expression of goal-tracking in goal-trackers, without affecting sign-tracking or ITI port entries. However, at a higher dose (0.1 mg/kg), dopamine D2-like receptor antagonism reduced sign-tracking, goal-tracking and ITI port entries.

CS+ lever activations (Figure 1d) differed by dose [F(2,42) = 9.982, *p* < 0.001] and phenotype [F(2,21) = 4.061, *p* = 0.032] with a significant dose × phenotype interaction [F(4,42) = 2.779, *p* = 0.039]. A dose of 0.1 mg/kg of eticlopride reduced CS+ lever activations in sign-trackers (0.1 mg/kg vs vehicle, *p* < 0.001; 0.1 mg/kg vs 0.01 mg/kg *p* = 0.001). However, 0.01 mg/kg of eticlopride had no effect on CS+ lever activations in sign-trackers, goal-trackers or intermediates (all *p*’s = 1).

ΔCS+ port entries (Figure 1e) were affected by eticlopride dose [F(2,42) = 13.676, *p* < 0.001] and phenotype [F(2,21) = 12.799, *p* < 0.001], with a significant dose × phenotype interaction [F(4,42) = 7.104, *p* < 0.001]. Compared to vehicle, ΔCS+ port entries were significantly lower in goal-trackers following 0.01 mg/kg (*p* = 0.005) or 0.1 mg/kg (*p* < 0.001) eticlopride, with no further reduction from 0.1 mg/kg to 0.01 mg/kg (*p* = 0.216). Intermediates made fewer ΔCS+ port entries following 0.1 mg/kg eticlopride compared to vehicle (*p* = 0.026), but not 0.01 mg/kg (*p* = 0.445).

Eticlopride reduced ITI port entries (Figure 1f), but only at the highest tested dose (0.1 mg/kg). There was no effect of phenotype [F(2,21) = 0.26, *p* = 0.774] or dose × phenotype interaction [F(2.691,28.256) = 0.216, *p* = 0.866]. However, there was an effect of dose [F(1.346,28.256) = 8.6, *p* = 0.003, ε= 0.673]. While 0.01 mg/kg did not differ from vehicle (*p* = 0.608), 0.1 mg/kg eticlopride reduced ITI port entries compared to vehicle (*p* = 0.014) and 0.01 mg/kg (*p* = 0.023). Effects on the CS− and Post-CS periods are shown in Figure A.4 and effects on probability of and latency to CS+ lever activations and port entries and PCA score are presented in Figure A.5, with statistical results in Tables A.4 and A.5, respectively.

#### Locomotor activity test in an open field arena

After habituation to the locomotor chambers, during which no differences between phenotypes were observed (Figure A.6), rats were randomly allocated to receive vehicle, 0.01 mg/kg or 0.1 mg/kg SCH-23390 in a between-subjects design. Compared to vehicle, SCH-23390 significantly reduced locomotor activity at both the high (0.1 mg/kg) and low (0.01 mg/kg) dose (n = 8/dose), and effects were milder at the lower dose. A Kruskal-Wallace test found an effect of dose [H(2) = 18.24, *p* < 0.001] on total distance travelled (Figure 2a). SCH-23390 reduced total distance travelled at 0.1 mg/kg (*p* < 0.001 vs vehicle; *p* = 0.033 vs 0.01 mg/kg), but not at 0.01 mg/kg (*p* = 0.269 vs vehicle). SCH-23390 also dose-dependently reduced the total number of movement episodes [Figure 2b; one-way ANOVA, F(2,21) = 154.939, *p* < 0.001]. The number of movement episodes was reduced, relative to vehicle, at both doses (0.01 mg/kg, *p* = 0.011; 0.1 mg/kg, *p* < 0.001). The 0.1 mg/kg dose further reduced movement episodes compared to 0.01 mg/kg (*p* < 0.001). SCH-23390 also increased time spent in the center of the open field arena [Figure 2c; H(2) = 16.819, *p* < 0.001]. The 0.1 mg/kg dose significantly increased center time (*p* = 0.008 vs vehicle; *p* < 0.001 vs 0.01 mg/kg), but 0.01 mg/kg had no effect (*p* = 1 vs vehicle).

**Figure 2.**
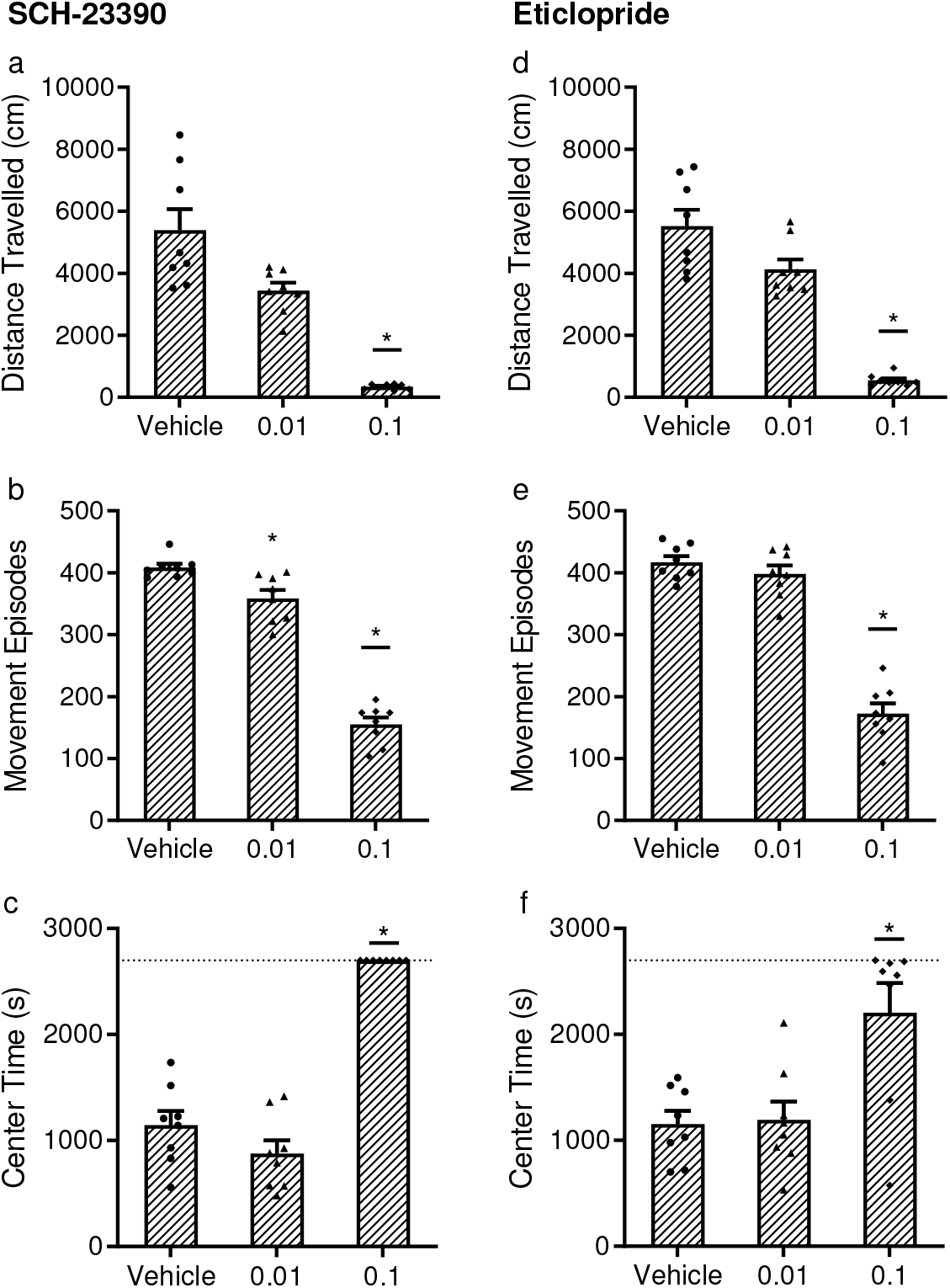
A reduction in locomotor behaviour by dopamine receptor antagonists. (a) Rats were randomly assigned to receive vehicle, 0.01, or 0.1 mg/kg of SCH-23390 prior to a 45-min locomotor test in an open field arena (n = 8/dose). Total distance travelled was significantly reduced at the 0.1 mg/kg but not 0.01 mg/kg dose. (b) The total number of movement episodes was reduced by SCH-23390 at both the 0.01 and 0.1 mg/kg doses. (c) The amount of time spent in the center of the open field arena was significantly increased by 0.1 mg/kg SCH-23390, reflecting rats’ near-immobility at this dose and the fact that all rats remained in the center for the entire session (2,700 s; dotted line). (d) On a separate testing day, eticlopride was tested using a similar design (n = 8/dose, randomly assigned). Eticlopride reduced total distance travelled at the 0.1 mg/kg but not 0.01 mg/kg dose. (e) Similarly, eticlopride reduced the total number of movement episodes at the 0.1 mg/kg but not 0.01 mg/kg dose. (f) Eticlopride only significantly increased center time at the 0.1 mg/kg dose. Data are means ± SEM. * *p* < 0.05 for Bonferroni corrected post-hoc comparisons.

Compared to vehicle, the high but not low dose of eticlopride reduced locomotor activity (n = 8/dose). Eticlopride reduced total distance travelled [Figure 2d; H(2) = 17.565, *p* < 0.001] at the 0.1 mg/kg dose (*p* < 0.001 vs vehicle; *p* = 0.024 vs 0.01 mg/kg), but not at 0.01 mg/kg (*p* = 0.413 vs vehicle). Eticlopride also reduced movement episodes [Figure 2e; one-way ANOVA, F(2,21) = 100.467, *p* < 0.001] at the 0.1 mg/kg dose (*p* < 0.001 vs vehicle and 0.01 mg/kg), but not at 0.01 mg/kg (*p* = 0.999 vs vehicle). Finally, eticlopride increased time spent in the center of the arena [Figure 2f; F(2,21) = 8.63, *p* = 0.002] at the 0.1 mg/kg dose (*p* = 0.004 vs vehicle; *p* = 0.006 vs 0.01 mg/kg), but not at 0.01 mg/kg (*p* = 1 vs vehicle).

### Effect of amphetamine-induced psychomotor sensitization on the acquisition of sign- and goal-tracking

#### Amphetamine exposure

Rats received either vehicle or 2 mg/kg amphetamine (n = 12/group) before exposure to the open field arena for 30 minutes (Figures 3a-c), followed immediately by placement into the conditioning chambers for 20 minutes (Figure 3d), in order to pair the effects of amphetamine with both contexts.

**Figure 3.**
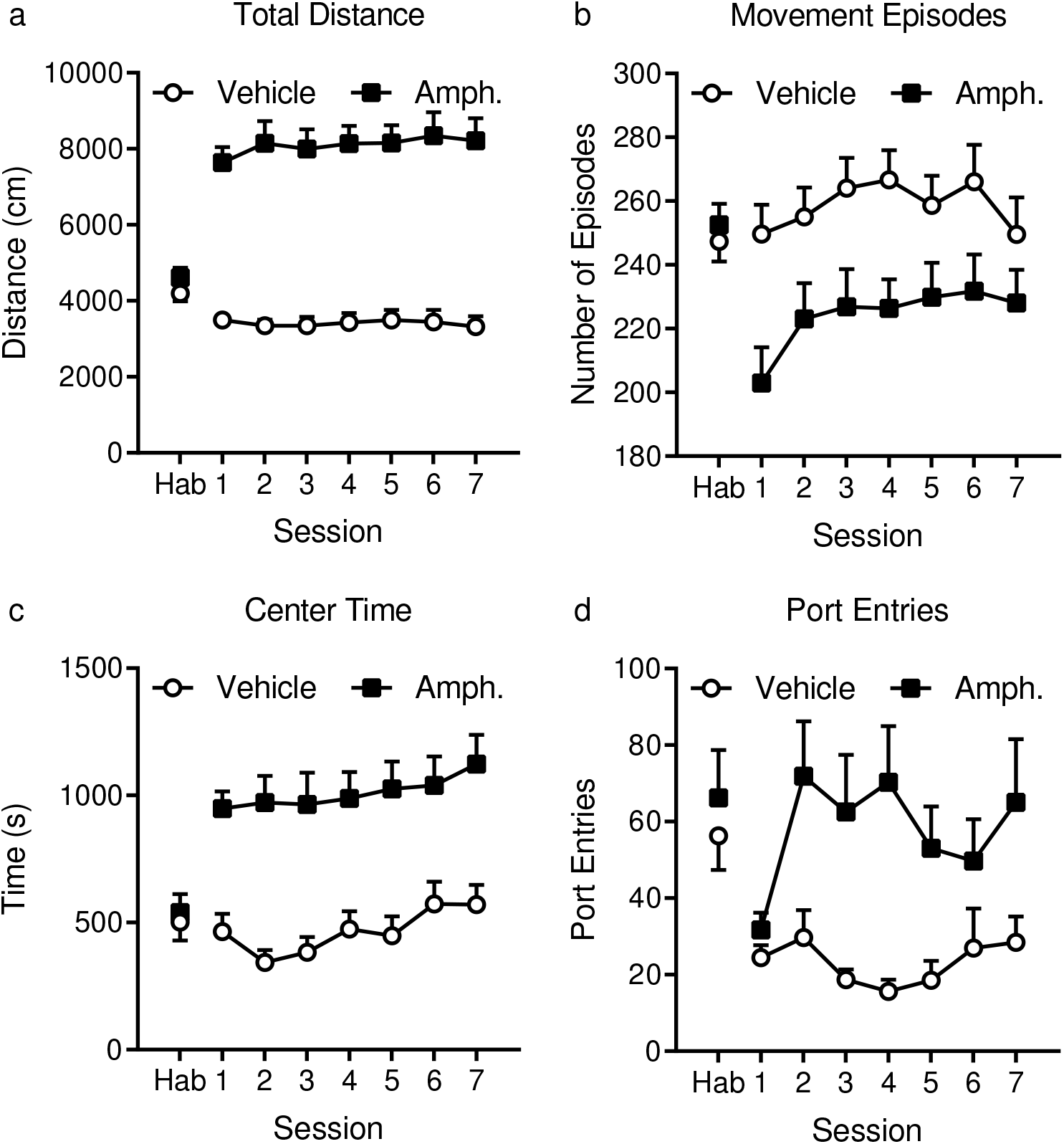
Repeated injections of amphetamine increased locomotor activity compared to vehicle. (a) Total distance travelled in 30 min was not significantly different between vehicle and amphetamine rats (n = 12/group) during a habituation session when all rats received saline. However, daily, 2 mg/kg amphetamine injections significantly increased total distance travelled. (b) Similarly, the number of movement episodes was not different during habituation, but was significantly reduced by amphetamine. (c) The amount of center time was not different between vehicle and amphetamine rats during habituation, but was increased by amphetamine. (d) Rats were immediately transferred to conditioning chambers after their locomotor session for 20 min. During habituation, there was no difference in the number of port entries made, but amphetamine rats made significantly more port entries during the amphetamine exposure phase. Data are means ± SEM.

During an initial, drug-free habituation session, there were no significant differences between groups in total distance travelled [Figure 3a; t(22) = −1.238, *p* = 0.229], number of movement episodes [Figure 3b; t(22) = −0.565, *p* = 0.578], time spent in the center of the arena [Figure 3c; t(22) = −0.365, *p* = 0.719], or port entries [Figure 3d; equal variances not assumed; t(19.888) = −0.646, *p* = 0.526].

Amphetamine treatment increased the total distance travelled [Figure 3a; F(1,22) = 89.115, *p* < 0.001], but there was no effect of session [F(3.51,77.214) = 0.438, *p* = 0.757, ε= 0.585] or session × treatment interaction [F(3.51,77.214) = 0.635, *p* = 0.619]. Amphetamine treatment decreased the number of movement episodes [Figure 3b; F(1,22) = 9.763, *p* = 0.005], suggesting increased movement per episode consistent with amphetamine’s psychostimulant effects. However, there was no effect of session [F(3.083,67.816) = 2.035, *p* = 0.115, ε= 0.514], or session × treatment interaction [F(3.083,67.816) = 0.584, *p* = 0.632]. Amphetamine treatment increased the amount of time spent in the center of the arena [Figure 3c; F(1,22) = 24.693, *p* < 0.001]. There was an effect of session [F(3.615,79.527) = 3.582, *p* = 0.012, ε= 0.602], but no differences between specific sessions or session × treatment interaction [F(3.615,79.527) = 0.655, *p* = 0.61]. Finally, amphetamine treatment was still effective in increasing psychomotor activity in the conditioning chamber, because amphetamine elevated the number of port entries made during the 20-min session [Figure 3d; F(1,22) = 11.878, *p* = 0.002] in the conditioning chamber, but there was no effect of session [F(3.442,75.716) = 1.801, *p* = 0.147, ε= 0.574] or session × treatment interaction [F(3.442,75.716) = 1.949, *p* = 0.121].

#### Acquistion of sign-tracking

Based on PCA scores from session 15 and 16, vehicle rats were mostly sign-trackers (n = 10) with 2 goal-trackers. Amphetamine rats were also mostly sign-trackers (n = 9) with 1 goal-tracker and 2 intermediates.

CS+ lever activations increased as a function of session [Figure 4a; F(3.202,70.437) = 11.309, *p* < 0.001, ε= 0.213], but there was no effect of prior amphetamine exposure on the acquisition of this behaviour [treatment: F(1,22) = 0.226, *p* = 0.639]. There was also no session × treatment interaction [F(3.202,70.437) = 1.024, *p* = 0.391]. For ΔCS+ port entries (Figure 4b), there was no effect of treatment [F(1,22) = 1.593, *p* = 0.22], session [F(2.597,57.13) = 2.49, *p* = 0.078, ε= 0.173], or session × treatment interaction [F(2.597,57.13) = 0.332, *p* = 0.773]. Similarly, there were no effects of prior amphetamine exposure on CS− lever activations, ΔCS− port entries, or Post-CS port entries (Figure A.7; Table A.6). While PCA scores (Figure 4c; see Figure A.8 for its components) shifted towards sign-tracking over 16 sessions [F(3.028,66.624) = 8.259, *p* < 0.001, ε= 0.202], there was no effect of treatment [F(1,22) = 0.332, *p* = 0.57] or session × treatment interaction [F(3.028,66.624) = 0.4, *p* = 0.756]. Thus, contrary to our hypothesis, prior, repeated amphetamine exposure had no effect on the development of sign-tracking and goal-tracking phenotypes over the course of training.

**Figure 4.**
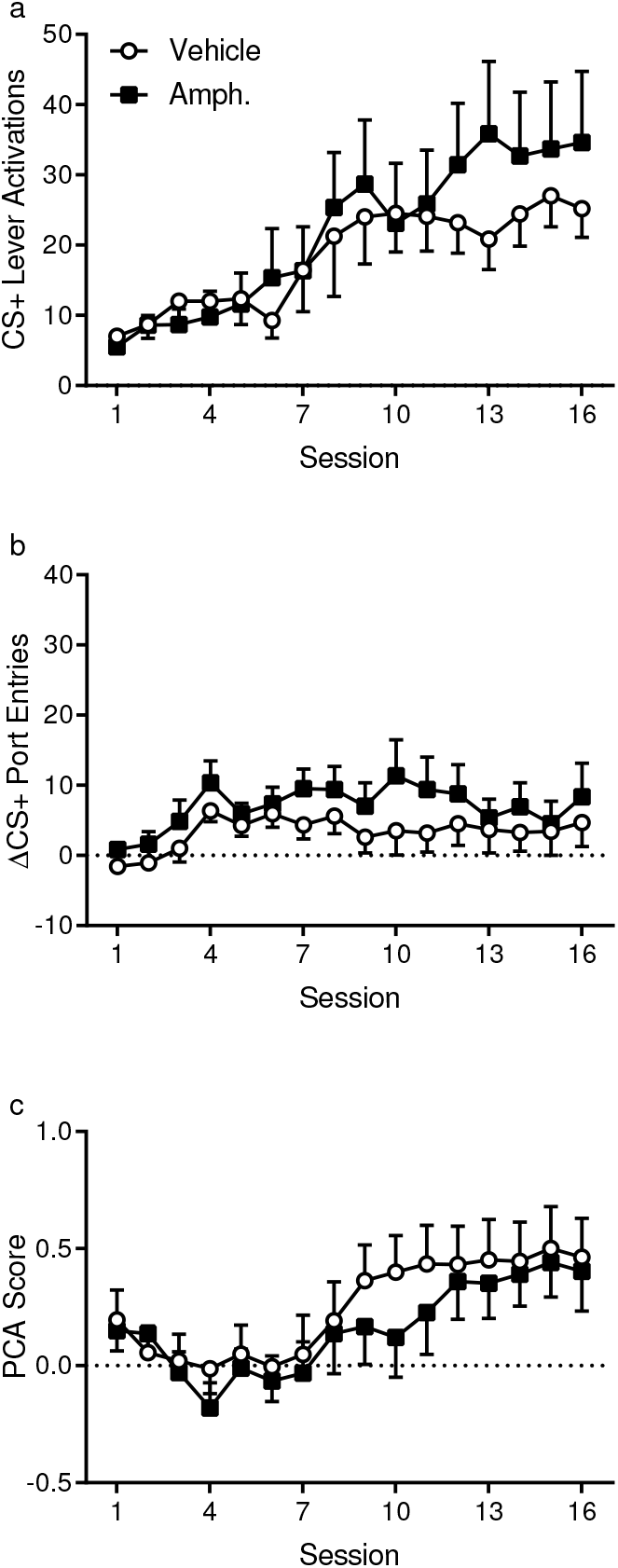
Acquisition of sign-tracking and goal-tracking did not differ between vehicle and amphetamine-exposed rats. (a) CS+ lever activations did not differ between rats that received vehicle or 2 mg/kg amphetamine during the amphetamine exposure phase (n = 12/group). (b) ΔCS+ port entries (CS+ port entries – Pre-CS+ port entries) also did not differ between groups during acquisition. (c) PCA scores showed that both cohorts acquired a sign-tracking phenotype overall and there were no significant differences in PCA scores during training. Data are means ± SEM.

#### Sensitization test

After receiving an acute 0.75 mg/kg amphetamine challenge, total distance travelled (Figure 5a) did not differ between vehicle and amphetamine-treated rats [t(22) = −1.121, *p* = 0.274]. However, amphetamine-treated rats initiated fewer movement episodes [Figure 5b; t(22) = 2.783, *p* = 0.011], suggesting increased movement per episode. Amphetamine-treated rats spent more time in the center of the arena [Figure 5c; t(22) = −2.179, *p* = 0.04]. Further exploration of the data showed that amphetamine-treated rats spent less time in the corners of the arena [Figure 5d; t(22) = 3.295, *p* = 0.003] and showed more locomotor activity in the center of the arena [Figure 5e; t(22) = −2.918, *p* = 0.008], indicating higher levels of exploratory activity than vehicle-treated rats. There was no evidence of an increase in stereotypy because rats in both groups spent, on average, less than two minutes engaged in restricted movements suggestive of stereotypic behaviours during the 30-min test session [Figure 5f; t(22) = 0.574, *p* = 0.572].

**Figure 5.**
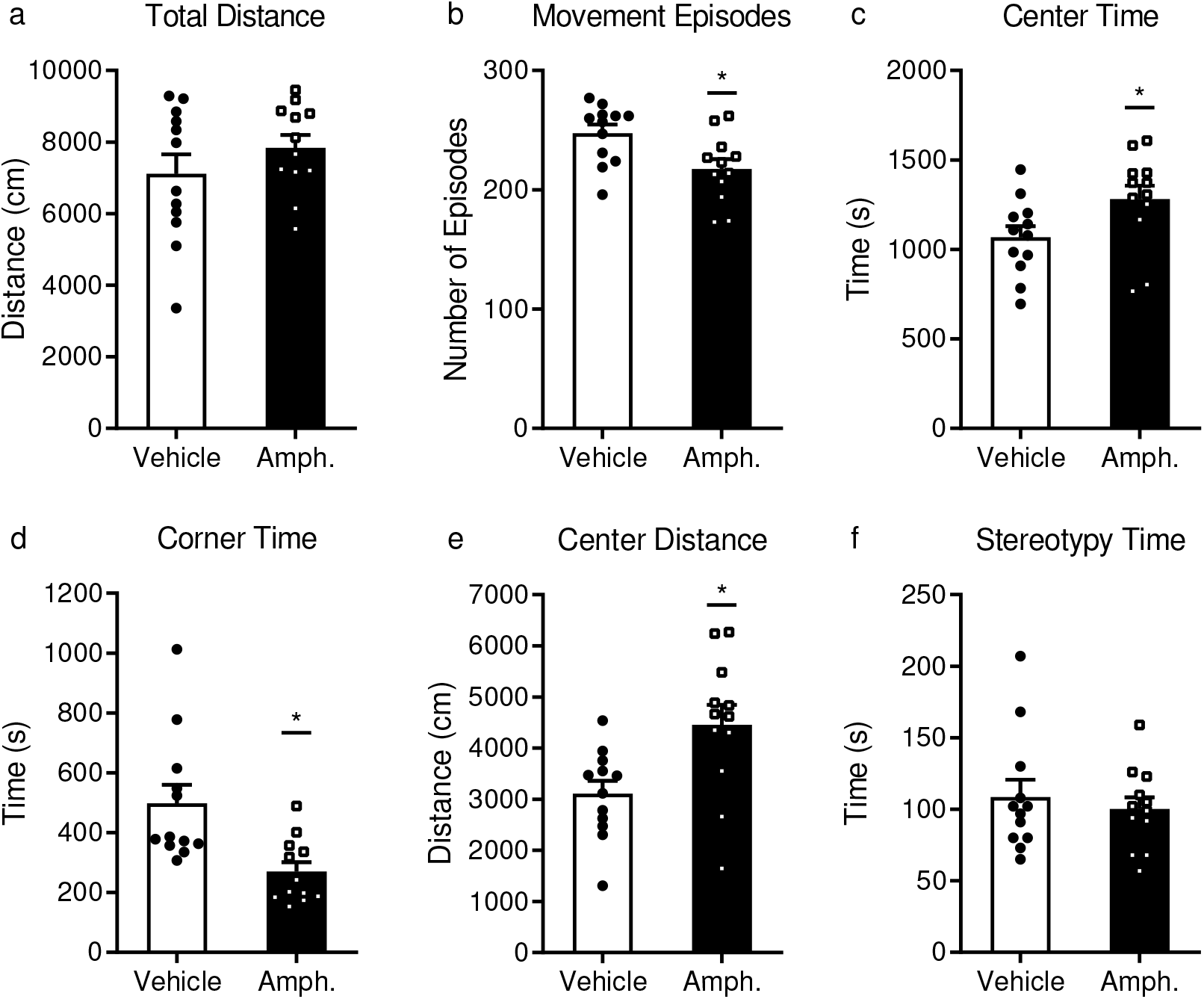
Amphetamine-exposed rats showed evidence of sensitization. Both vehicle and amphetamine-exposed rats (n = 12/group) received a 0.75 mg/kg amphetamine challenge dose before a locomotor activity test in an open field arena. (a) Although total distance did not significantly differ between groups during the 30 min test, (b) amphetamine-exposed rats had fewer movement episodes than vehicle rats, suggesting more movement per episode, and (c) spent more time in the center of the open field arena than vehicle rats. Further exploration of the data showed that (d) amphetamine-exposed rats spent less time in the corners of the arena and (e) travelled a greater distance in the center of the arena. (f) Time spent in restricted movements suggestive of stereotypic behaviours was low in both vehicle and amphetamine-exposed rats. Data are means ± SEM. * *p* < 0.05 for an independent t-test.

## Discussion

We found that antagonism of dopamine D1-like receptors using SCH 23390 following 20 sessions of PCA training reduced the expression of sign-tracking and goal-tracking conditioned responses when tested under extinction conditions. However, SCH 23390 also reduced ITI port entries and locomotor activity in an open field arena. The D2-like receptor antagonist, eticlopride, reduced sign- and goal-tracking behaviour, ITI port entries and locomotor activity at the 0.1 mg/kg dose. However, 0.01 mg/kg of eticlopride selectively reduced goal-tracking behaviour in goal-trackers, without affecting sign-trackers or intermediates, and this dose did not reduce ITI port entries or locomotor activity in an open field arena. We also found that psychomotor sensitization produced by repeated amphetamine pre-treatment did not alter the subsequent acquisition of sign-tracking behaviour. These results suggest that after extended PCA training, dopaminergic activity at D2 receptors is important for the expression of goal-tracking, but not for the expression of sign-tracking.

Following extended PCA training in Experiment 1, D1 antagonism using SCH-23390 disrupted both sign-tracking and goal-tracking; however, the interpretation of these effects is confounded by data suggesting that SCH 23390 pre-treatment produced a motor suppressive effect. We chose to test 0.01 mg/kg SCH-23390 because this dose impaired Pavlovian conditioned responding elicited by an auditory alcohol-predictive CS without affecting ITI port entries (Sciascia et al., 2014; Sparks et al., 2014). Other laboratories have shown that this dose disrupted the acquisition of sign-tracking (Chow et al., 2016) and expression of sign-tracking (Clark et al., 2013). Moreover, 0.01 mg/kg SCH-23390 has been shown to increase cocaine self-administration (Koob et al., 1987), reduce saccharin seeking (Aoyama et al., 2016) and alter risky decision-making in rats (Smith et al., 2018). Although our previous studies did not find effects of SCH-23390 on ITI port entries (Sciascia et al., 2014) and other laboratories have reported no effect of SCH-23390 on food consumption during Pavlovian conditioning (Chow et al., 2016) or on Pre-CS locomotor activity (Palmatier et al., 2014), we found that 0.01 mg/kg SCH-23390 impaired ITI port entries and the number of movement episodes in a locomotor activity test. This dose is even lower than doses that produced non-specific effects reported by Smith et al. (2018), who observed that rats stopped eating pellets after 0.017 and 0.03 mg/kg. While SCH-23390 appeared to reduce sign-tracking in sign-trackers and goal-tracking in goal-trackers, it is difficult to interpret these effects in the presence of the observed motor-suppressing effects. Moreover, the appearance of a phenotype-specific effect may be due to floor effects because sign-trackers do not make many port entries and goal-trackers do not make many lever responses. Therefore D1-like receptor antagonism may have effects on sign-tracking or goal-tracking, but further studies are required using lower doses of a D1 antagonist that do not produce motor suppressive effects.

We found that following extended PCA training, D2 antagonism using eticlopride reduced goal-tracking in goal-trackers, but not sign-tracking in sign-trackers, at a dose that did not affect ITI responses. Some studies have found that D2 antagonists administered prior to testing reduced both sign-tracking and goal-tracking (Fraser et al., 2016; Lopez et al., 2015), which might initially appear inconsistent with the present findings that sign-tracking was unaffected. However, these differences may be due to different testing parameters, such as when antagonists were administered or testing conditions. When administered intermittently prior to training, eticlopride reduced both sign-tracking and goal-tracking responses, but left sign-tracking, goal-tracking, and the conditioned reinforcing effects of the CS lever largely intact during drug-free tests (Chow et al., 2016). Chow et al. (2016) suggest that this may be because D2-like receptors are important for the performance of these behaviours, but not for learning the CS-US association. Importantly, we tested the effect of eticlopride under extinction conditions (no sucrose), similar to the drug-free CS only tests used by Chow et al. (2016). In contrast, studies that have observed effects on both sign-trackers and goal-trackers did not test under extinction conditions (Fraser et al., 2016; Lopez et al., 2015). Testing under extinction conditions ensured that any observed effects would relate specifically to performance of already-learned behaviours and would not be influenced by the reinforcer. Consistent with the interpretation of Chow et al. (2016), Roughley & Killcross (2019) observed an impairment of sign-tracking, but not goal-tracking, in drug-free sessions that followed one week of daily pre-training eticlopride. Our study also tested the effect of eticlopride, but administered immediately prior to testing under extinction conditions rather than during training. The present data therefore extend these findings, by showing that D2 antagonism prior to testing under extinction conditions disrupts goal-tracking, which is entirely reliant on the CS-US association but leaves sign-tracking, which involves phasic dopamine responses to the CS rather than to the US (Flagel et al., 2011), intact. Moreover, our findings are consistent with previous studies showing that treatment with the antipsychotic/D2-like receptor antagonist, haloperidol, did not affect the expression of established sign-tracking (Bédard et al., 2011). Some caution is required in interpreting our finding that D2 antagonism disrupts goal-tracking under extinction, because although locomotor behaviour in the open field arena was not significantly reduced, there may have been a slight reduction in distance travelled at the 0.01 mg/kg dose. Testing under other parameters, such as during reinforced sessions, may also alter the results and implications. Therefore, our results warrant further investigation with lower systemic doses or using intracranial manipulations to more confidently dissociate the effects on PCA behaviour from motor suppressive effects, under a variety of experimental parameters.

One limitation of our design is that we always tested rats with eticlopride after they had received tests with SCH-23390. We did all of our tests within-subjects to minimise the number of rats used and maximise statistical power, however, if administration of SCH-23390 had long-term effects on PCA behaviour it could confound the eticlopride results. However, we did provide a day of normal PCA training between drug tests to allow rats to return to baseline and dopamine receptor supersensitivity is typically associated with chronic, continuous administration of dopamine antagonists in rats (Ericson et al., 1996; Samaha et al., 2008; Servonnet & Samaha, 2020), in contrast to the acute injections used here. There was also no significant difference in performance in the vehicle tests for SCH-23390 and eticlopride with respect to CS+ or CS− lever activations, ΔCS+ or ΔCS− port entries and ITI port entries (paired t-tests, *p* > 0.05). Therefore, a confound from drug testing order is not obvious in our data, but remains a possibility.

Another consideration is that the rats in our study were skewed towards sign-tracking. In Experiment 1, 14 of the 24 rats were classified as sign-trackers (58%), 6 rats were goal-trackers (25%) and 4 were intermediates (17%). In comparison, previous work from our laboratory that analysed results across 5 experiments found a more even distribution with approximately 32 of 76 rats (42%) that were sign-trackers, while 28 (37%) were goal-trackers (Villaruel & Chaudhri, 2016). Other laboratories have also found approximately even distributions of sign- and goal-tracking in Long-Evans rats. Lopez et al. (2015) reported the same number of sign- and goal-trackers after 4 sessions (9 of 45 rats each), with the remaining 27 (60%) of an intermediate phenotype. This may imply that the rats used in this study were predisposed towards sign-tracking, which may limit the generalisability of these results. Moreover, the small numbers of goal-trackers and intermediates here suggests caution is required because of low statistical power, although statistically significant effects for goal-trackers indicates that the effects of eticlopride are likely to be large. It appears then, that Long Evans rats are highly variable with respect to their PCA phenotypes and that future studies are required to replicate these results in cohorts with different predispositions towards sign-tracking.

In Experiment 2, we observed acquisition of sign-tracking with extended PCA training. This finding replicates our previous studies, where some rats shifted from goal-tracking to sign-tracking with an alcohol cue after 16 or more PCA sessions (Srey et al., 2015; Villaruel & Chaudhri, 2016). In the present study, which used a sucrose reinforcer, rats appeared to be acquiring a goal-tracking response in sessions 1-4 before PCA scores increased and reached asymptote around session 10-13. This result suggests a robust tendency (in our hands) for rats to eventually acquire a sign-tracking phenotype across reinforcers.

Contrary to our expectations, prior amphetamine exposure did not affect the acquisition of sign-tracking, despite several previous studies implicating dopamine signalling in the acquisition of sign-tracking. Sign-trackers show greater expression of D1 receptors after the first training session and dopamine signalling remains important for the maintenance of sign-tracking responses (Flagel et al., 2007; Fraser et al., 2016). Moreover, a higher phasic dopamine response was associated with sign-tracking across multiple sessions (Flagel et al., 2011). Previous studies found that psychomotor sensitization to amphetamine augmented CS lever activations (Wyvell & Berridge, 2001), and augmentation may be achieved with a single amphetamine injection (Schuweiler et al., 2018; Wyvell & Berridge, 2000). However, other studies have shown that amphetamine-induced psychomotor sensitization enhanced goal-tracking (Simon et al., 2008). One consideration is that these studies used very different protocols, such as inducing sensitization after conditioning (Wyvell & Berridge, 2001) or using food-restricted rats (Simon et al., 2008). However, it has also been shown that the dopamine response to an appetitive CS diminishes with extended training (Clark et al., 2013), suggesting that although dopamine is important for the maintenance of the sign-tracking response (Fraser et al., 2016), additional dopamine signalling may not be required to express a sign-tracking response. Thus, previous studies have shown that dopamine is required for expressing a sign-tracking response, and our data suggest that sensitization-related plasticity within the dopamine system is not sufficient to enhance the rate of acquisition of sign-tracking.

An alternative interpretation of this result is that because the majority of rats in our study became sign-trackers, a ceiling effect prevented any influence of amphetamine-induced sensitization-related plasticity on PCA behaviour from becoming apparent. Our study was designed to examine the rate at which rats progress to sign-tracking, which appears similar in both experimental groups and takes 7-10 days to become clear. However, another possible approach to this question is determining whether sensitization could alter the final distribution of phenotypes even if the rate of acquisition of those phenotypes is not affected. Addressing this would require testing rats that are less predisposed to sign-tracking than the ones used here. A related concern is that we gave amphetamine prior to PCA training, so it is unknown whether rats would have expressed their predominantly sign-tracking phenotypes regardless of amphetamine treatment. Although giving amphetamine before PCA training, with rats randomly allocated to experimental conditions, allowed us to examine any potential causal effects on the rate of progression to a sign-tracking phenotype, another approach for future studies is to use animals that are predisposed to goal-tracking and examine whether sensitization is able to accelerate acquisition of sign-tracking.

Data from our sensitization test, where all rats were given an amphetamine challenge, provides robust evidence that our amphetamine exposure regimen produced sensitization of locomotor behaviour across multiple measures of locomotor activity, even though this did not affect the acquisition of sign-tracking. We exposed rats first to the open field arena and then to the conditioning chambers during the amphetamine exposure regimen because sensitization is context-dependent (Badiani et al., 1995a,b; Crombag et al., 2001, 2000). Since brain concentrations of amphetamine peak within 5-30 min and decline with a t½ of 40-70 min (Coutts et al., 1986; Lokiec et al., 1978), our exposure timeline meant that rats had high brain concentrations of amphetamine in both contexts. Our sensitization test followed previous studies, administering a 0.75 mg/kg amphetamine challenge to all rats (Robinson et al., 2015). An enhanced response to an amphetamine challenge after repeated exposure to the drug is a reliable between-subjects index of the development of psychomotor sensitization, because it has previously been shown that sensitization can require weeks of abstinence from experimenter-administered amphetamine before it is expressed (Paulson & Robinson, 1995). In contrast to vehicle rats, rats that were previously exposed to amphetamine had fewer movement episodes, suggesting increased movement per episode, increased center time and decreased corner time, consistent with increased exploratory behaviour. While there was no difference in total distance travelled during the sensitization test, total distance travelled at test was similar to the exposure phase, suggesting a ceiling effect. Previous studies have found that differences in locomotor activity emerge later in the sensitization test (Simon et al., 2008), so an effect on total distance travelled may have emerged if our sensitization test had lasted longer. We also observed significantly more distance travelled in the center of the arena in amphetamine-treated rats. Along with low levels of restricted movements suggestive of stereotypies, the center distance travelled suggests rats were ambulatory, providing further evidence of enhanced locomotor activity. The effect of amphetamine on multiple locomotor activity measures during the sensitization test therefore strongly suggests that repeated amphetamine exposure induced locomotor sensitization, but that this did not alter the acquisition of sign-tracking behaviour.

It is possible that other methods for intervening in the dopamine system could still alter the acquisition of sign-tracking behaviour. For example, it has recently been shown that inhibition of prelimbic cortex projections to the paraventricular nucleus of the thalamus can shift goal-trackers towards sign-tracking (Campus et al., 2019). Although the prelimbic projections themselves are glutamatergic rather than dopaminergic, their inhibition was shown to increase dopamine levels in the nucleus accumbens via functional connectivity with other brain regions (Campus et al., 2019). It is therefore possible that more temporally or anatomically precise dopamine manipulations using opto- or chemogenetics may facilitate a shift to a sign-tracking phenotype. Nonetheless, our data suggest that increasing the overall sensitivity of the dopamine system through repeated psychostimulant administration does not predispose animals towards accelerated incentive salience acquisition.

Further studies are also required to determine the extent to which the results from the present studies generalise to work conducted under different conditions. For example, in the present study we used individually housed male rats and previous studies have shown that social isolation—at least in adolescence—can decrease baseline dopamine concentrations and enhance evoked dopamine release and uptake in the nucleus accumbens core and shell (Karkhanis et al., 2016). Prior work on dopamine regulation of sign-tracking and goal-tracking have used both individually and paired/group-housing conditions (Table 1). However, at present there appears to be too much variation in experimental procedures to draw firm conclusions about whether housing conditions strongly influence dopamine’s role in Pavlovian conditioned approach. Of note, previous studies have found that social housing does not alter Pavlovian conditioned approach in adult rats (Anderson et al., 2013). Further experiments are also required to optimise these procedures using female animals and test whether observed effects are sex-specific. The present work can guide future experiments on this issue. The importance of studying both sexes in this context is further highlighted by observations that i) previous studies, and the present one, on the role of dopamine in sign-tracking and goal-tracking have used male animals exclusively (Table 1), and ii) female rats can show higher levels of sign-tracking compared to males (Stringfield et al., 2019).

## Conclusion

The present studies sought to evaluate the role of the dopamine system in the sign- versus goal-tracking form of Pavlovian approach behaviour following extended PCA training. Antagonism of dopamine D1-like receptors reduced the expression of both sign- and goal-tracking conditioned responses, but these results should be interpreted with caution in the face of reduced ITI port entries and general locomotor behaviour. Interestingly, dopamine D2-like receptor signalling was specifically required for the expression of goal-tracking in goal-trackers, and this finding extends previous work on the effects of D2 receptor manipulations on conditioned approach behaviour (Bédard et al., 2011; Chow et al., 2016; Roughley & Killcross, 2019). A second, parallel objective was to investigate whether the dopamine system is involved in the shift from goal- to sign-tracking behaviour resulting from extended PCA training. Although we observed robust amphetamine-induced psychomotor sensitization on multiple measures of locomotor behaviour, it did not affect the rate at which rats shifted from goal-tracking to sign-tracking. However, it is possible that more temporally or anatomically precise dopaminergic manipulations could alter the trajectory of sign- and goal-tracking across extended PCA training. Further studies are therefore required to investigate the neurobiological mechanisms underlying the acquisition of sign-tracking following extended PCA training.

## Declarations

### Acknowledgements

This research was supported by grants from the Canadian Institutes of Health Research (CIHR; MOP-137030; NC), the Natural Sciences and Engineering Research Council (NSERC; RGPIN-2017-04802, NC), Fonds de la recherche du Québec – Santé (Chercheur-Boursier, NC) and Concordia University (Center for Studies in Behavioral Neurobiology, NC). SYK was supported by a Concordia Horizon Postdoctoral Fellowship and a postdoctoral fellowship from the Fonds de la recherche du Québec – Santé (FRQS). AU was supported by a Concordia Undergraduate Student Research Award. ANS was supported by the Fonds de la recherche du Québec – Santé (Chercheure-Boursiere). The authors gratefully acknowledge Uri Shalev for access to locomotor testing equipment.

### Author Contributions

SYK contributed to conceptualization, methodology, software, validation, formal analysis, investigation, data curation, writing – original draft, writing – review & editing, visualization and supervision of AU. AU contributed to conceptualization, methodology, formal analysis, investigation and writing – review & editing. ANS contributed to writing – review & editing. NC contributed to conceptualization, methodology, funding acquisition, resources, project administration, supervision of SYK and AU, visualization, writing – original draft and writing – review & editing.

## Appendix: Additional Figures and Tables

**Figure A.1.**
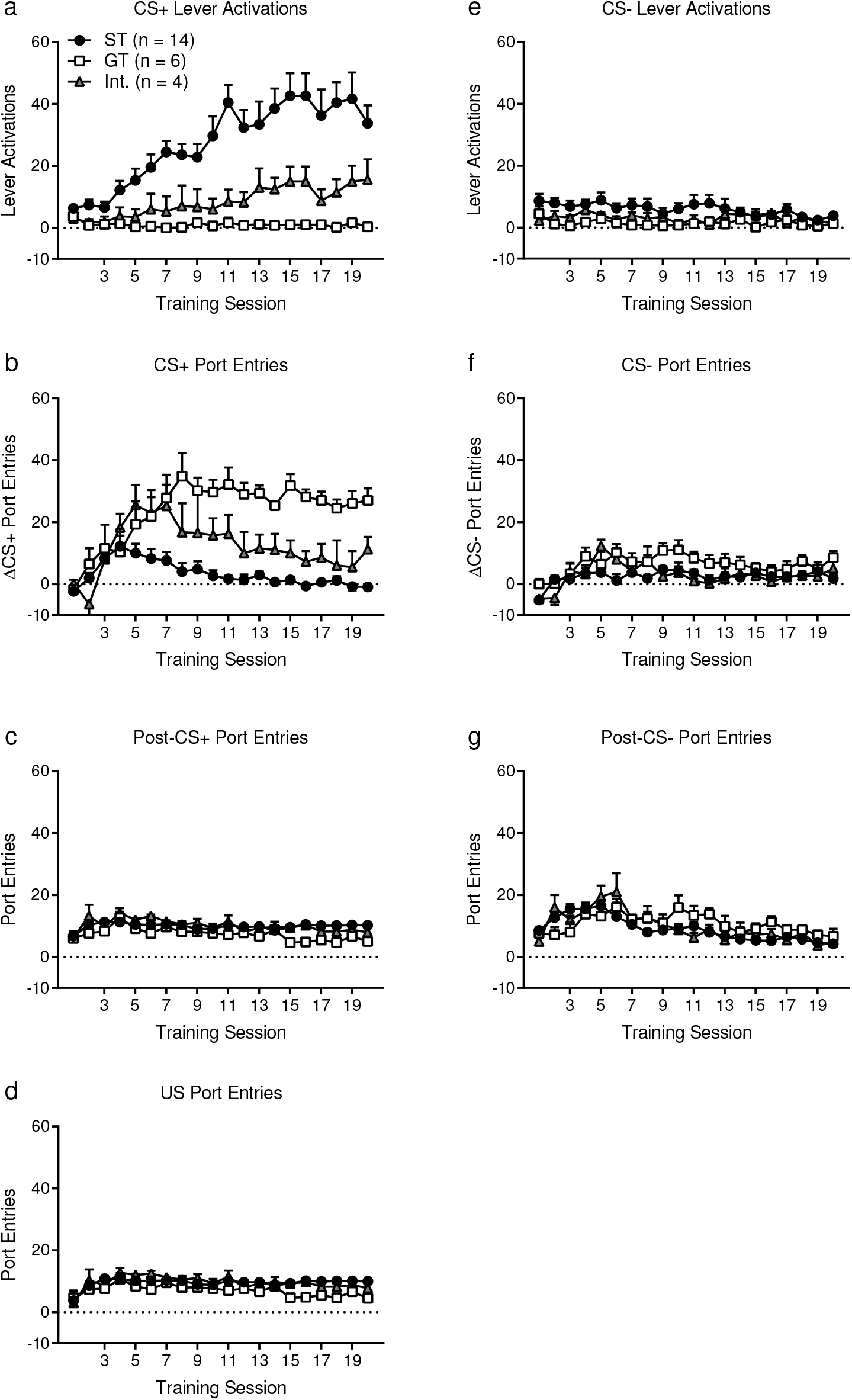
Sign-trackers and goal-trackers acquired distinct behaviours during Pavlovian PCA training sessions. (a) Rats classified as sign-trackers (ST; n = 14) interacted most with a tactile lever cue that predicted 10% sucrose delivery (CS+), while goal-trackers (GT; n = 6) and intermediates (Int.; n = 4) produced fewer CS+ lever activations. (b) ΔCS+ port entries (CS+ port entries minus Pre-CS+ port entries) were highest in goal-trackers. (c) Post-CS+ port entries were consistent throughout PCA training. (d) US port entries were made during the first 6 s of the Post-CS+ during operation of the syringe pump. Over 90% of Post-CS+ port entries were made during the US. (e) The overall level of interaction with a non-predictive lever cue (CS−) remained low throughout 20 sessions of training. (f) ΔCS− port entries (CS− port entries minus Pre-CS− port entries) remained low throughout training. (g) Post-CS− port entries also remained low throughout training. Each cue lever was available for 10 trials for 10 s per trial (total of 100 s/session). Data are means ± SEM. See Table A.1 for statistical results.

**Figure A.2.**
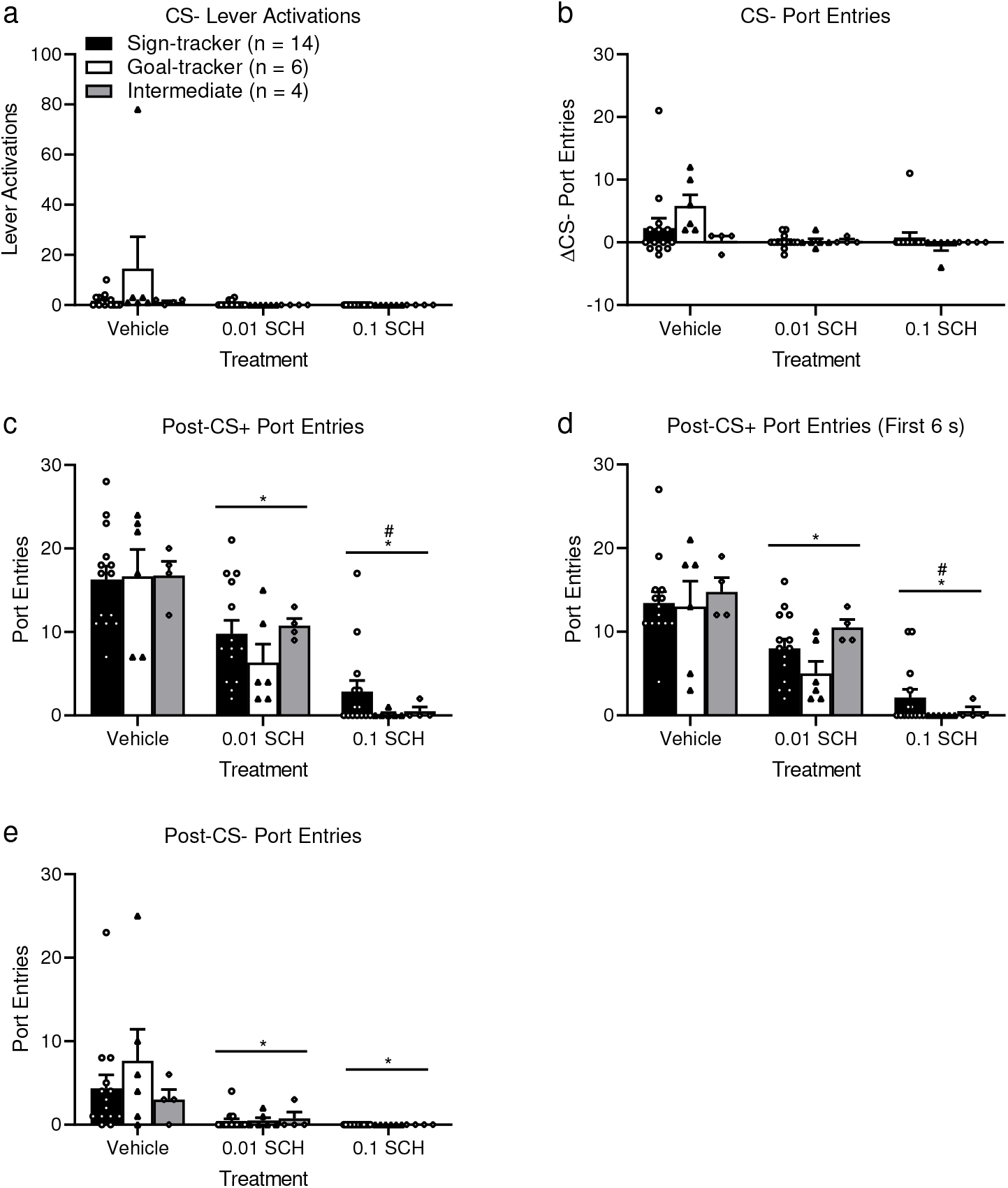
Effects of SCH-23390 on CS− responses and Post-CS responding. (a) SCH-23390 had no effect on CS− lever activations and (b) ΔCS− port entries. Both 0.01 mg/kg and 0.1 mg/kg SCH-23390 reduced port entries during (c) the full 10 s of the Post-CS+ period, (d) the first 6 s of the Post-CS+ period which corresponds to the operation of the syringe pump (which was empty at test) and (e) Post-CS− port entries. Data are means ± SEM. * *p* < 0.05 vs Vehicle; # *p* < 0.05 vs 0.01 SCH. See Table A.2 for statistical results.

**Figure A.3.**
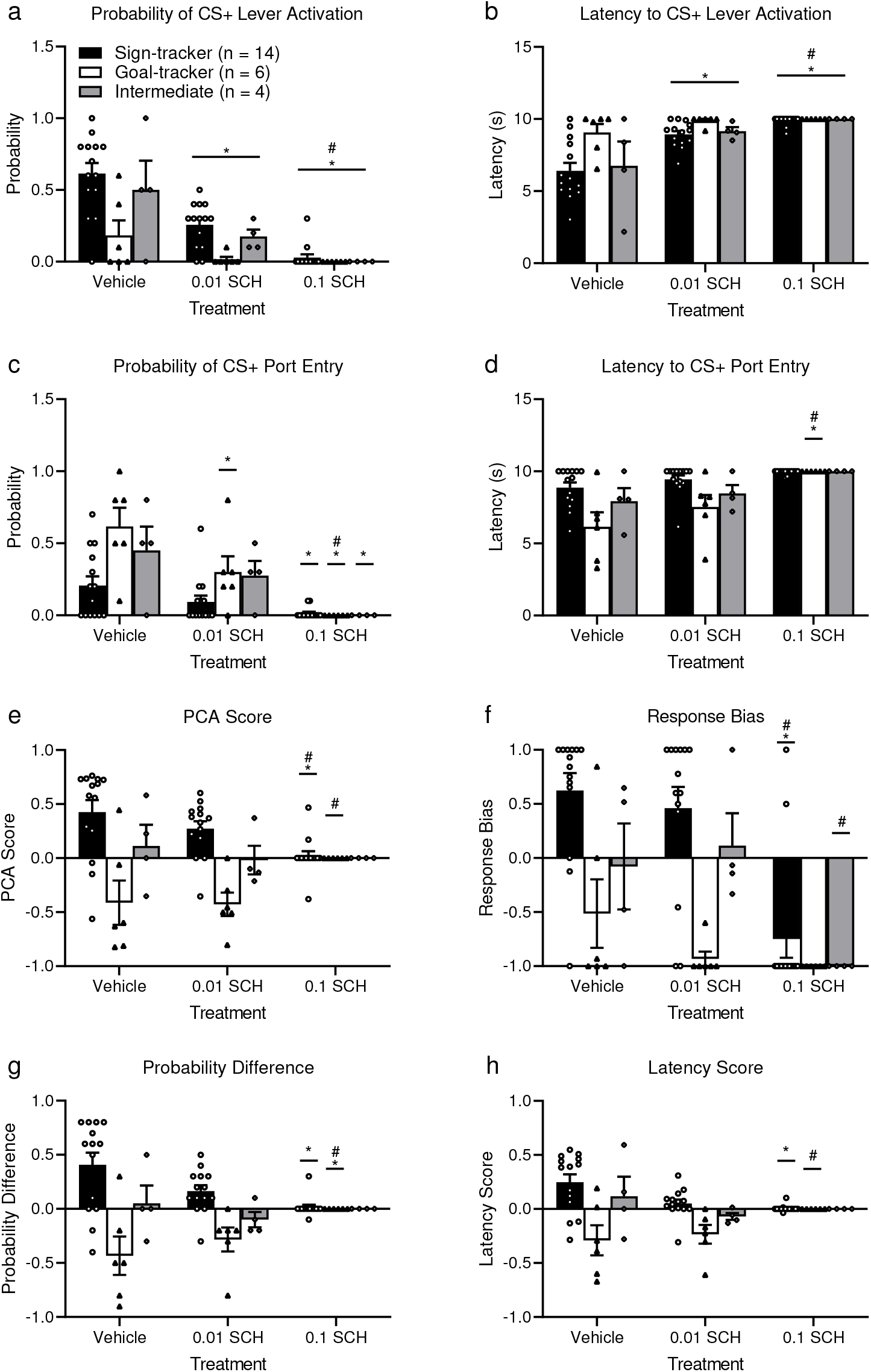
Effects on SCH-23390 on response probability, latency and PCA Score. (a) Probability of CS+ lever activation per trial, (b) latency to CS+ lever activation, (c) probability of CS+ port entries per trial, (d) latency to CS+ port entry. (e) PCA score and its components measures: (f) response bias, (g) probability difference and (h) latency score. Data are means ± SEM. * *p* < 0.05 vs Vehicle; # *p* < 0.05 vs 0.01 SCH. See Table A.3 for statistical results.

**Figure A.4.**
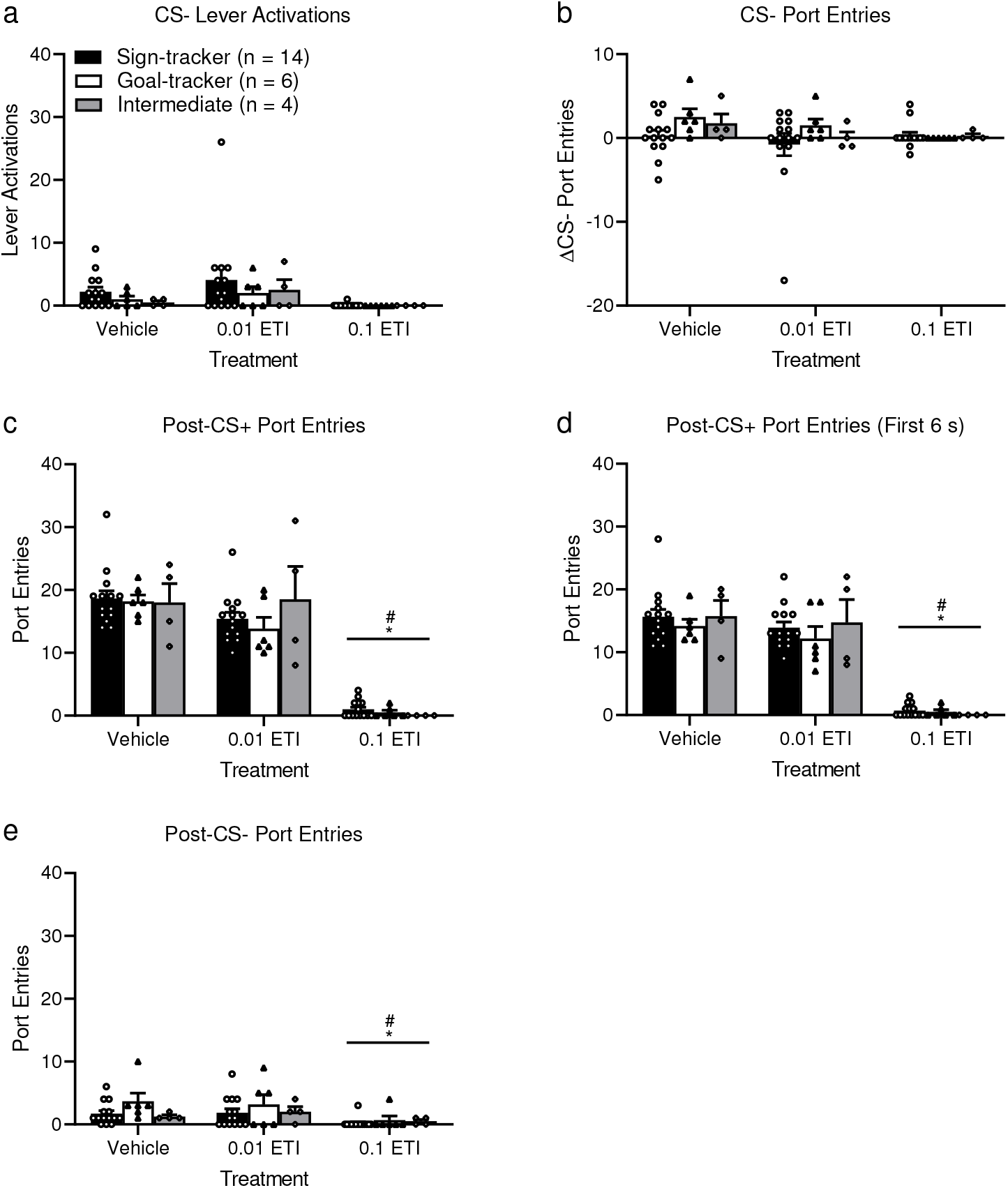
Effects of eticlopride on CS− responses and Post-CS responding. (a) Eticlopride had no effect on CS− lever activations and (b) ΔCS− port entries. Only 0.1 mg/kg eticlopride reduced port entries during (c) the full 10 s of the Post-CS+ period, (d) the first 6 s of the Post-CS+ period which corresponds to the operation of the syringe pump (which was empty at test) and (e) Post-CS− port entries. Data are means ± SEM. * *p* < 0.05 vs Vehicle; # *p* < 0.05 vs 0.01 ETI. See Table A.4 for statistical results.

**Figure A.5.**
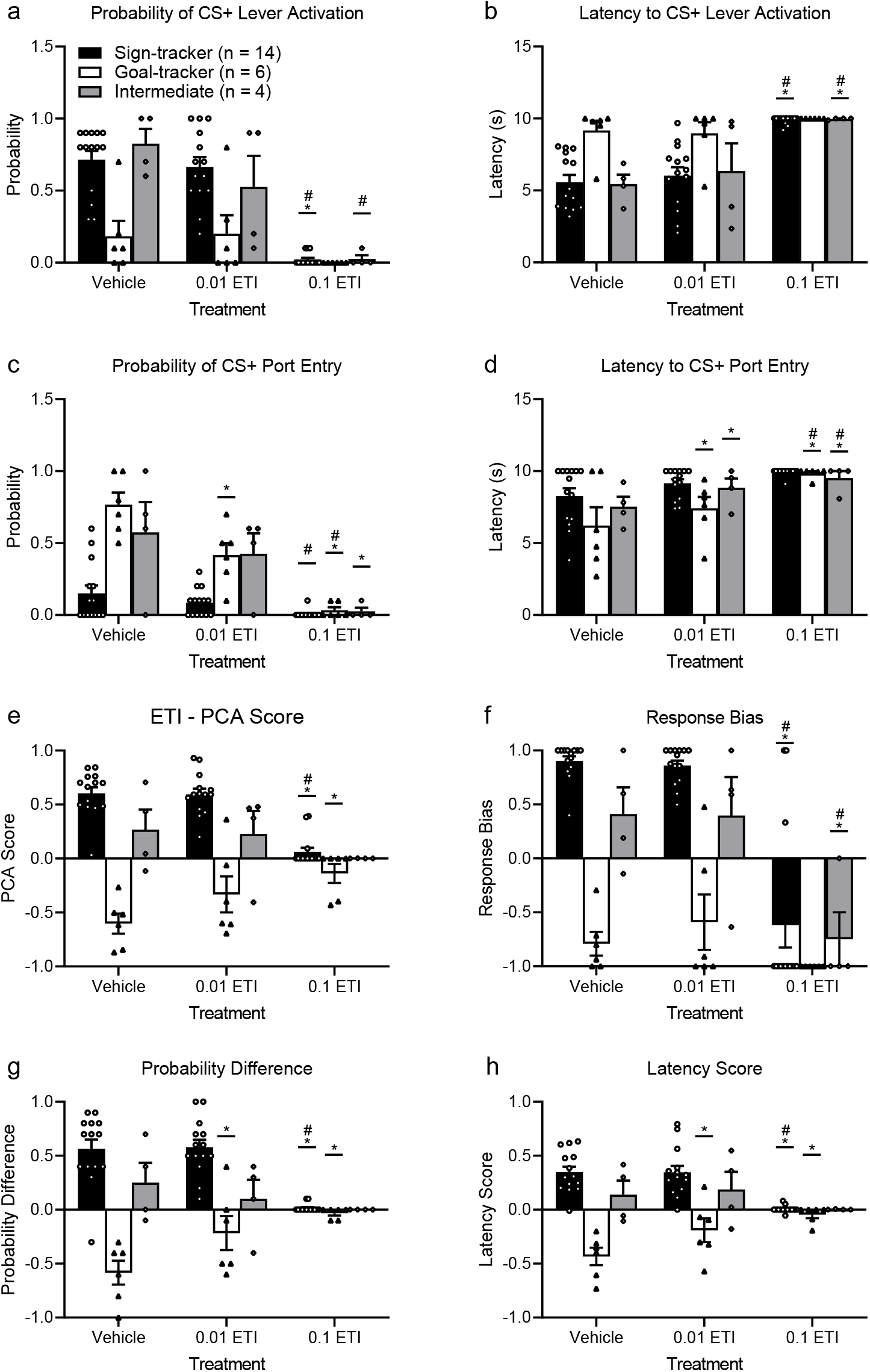
Effects on eticlopride on response probability, latency and PCA Score. (a) Probability of CS+ lever activation per trial, (b) latency to CS+ lever activation, (c) probability of CS+ port entries per trial, (d) latency to CS+ port entry. (e) PCA score and its components measures: (f) response bias, (g) probability difference and (h) latency score. Data are means ± SEM. * *p* < 0.05 vs Vehicle; # *p* < 0.05 vs 0.01 ETI. See Table A.5 for statistical results.

**Figure A.6.**
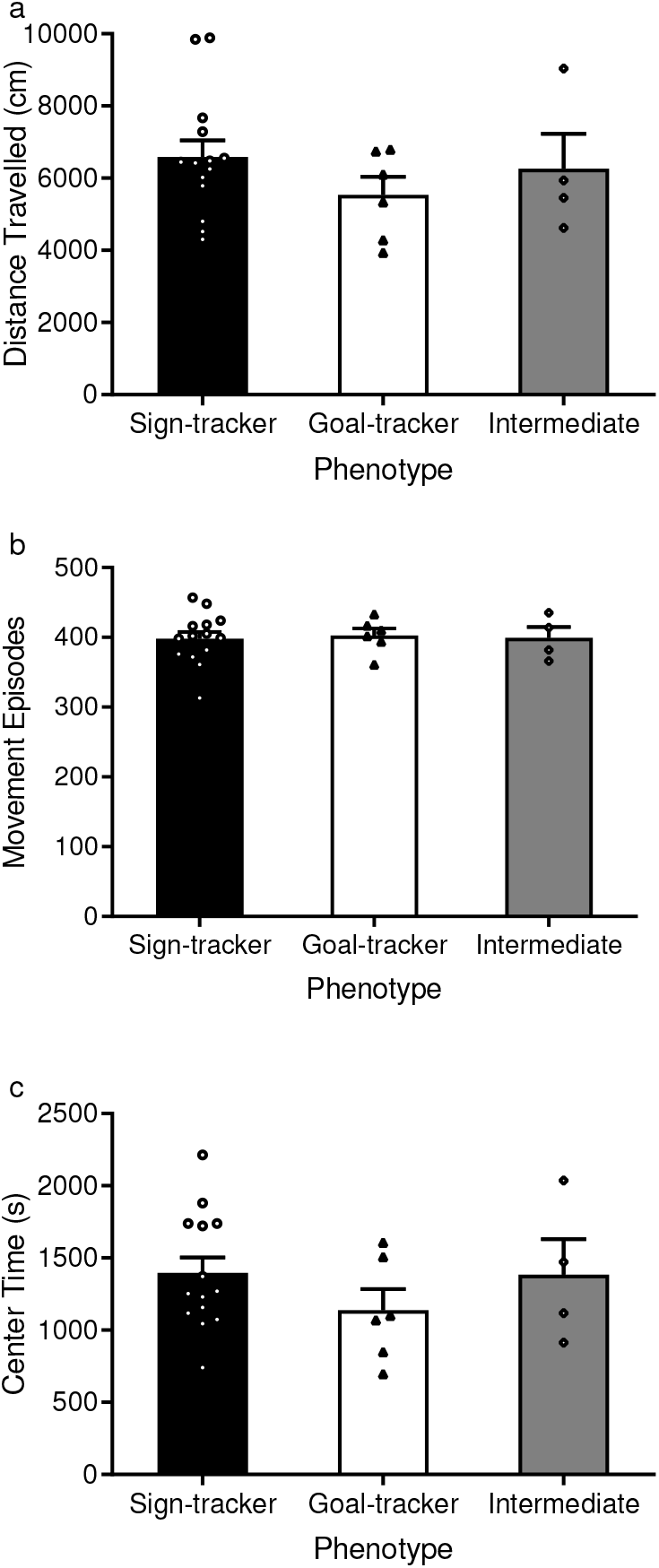
Locomotor behaviour during the habituation session did not differ by phenotype. Pavlovian conditioned approach phenotype (sign-tracker, goal-tracker or intermediate) did not affect (a) distance travelled [One-way ANOVA: F(2,21) = 0.883, *p* = 0.428], (b) the number of movement episodes [One-way ANOVA: F(2,21) = 0.045, *p* = 0.956] or (c) time spent in the center of the arena [One-way ANOVA: F(2,21) = 0.891, *p* = 0.425]. Data are means ± SEM.

**Figure A.7.**
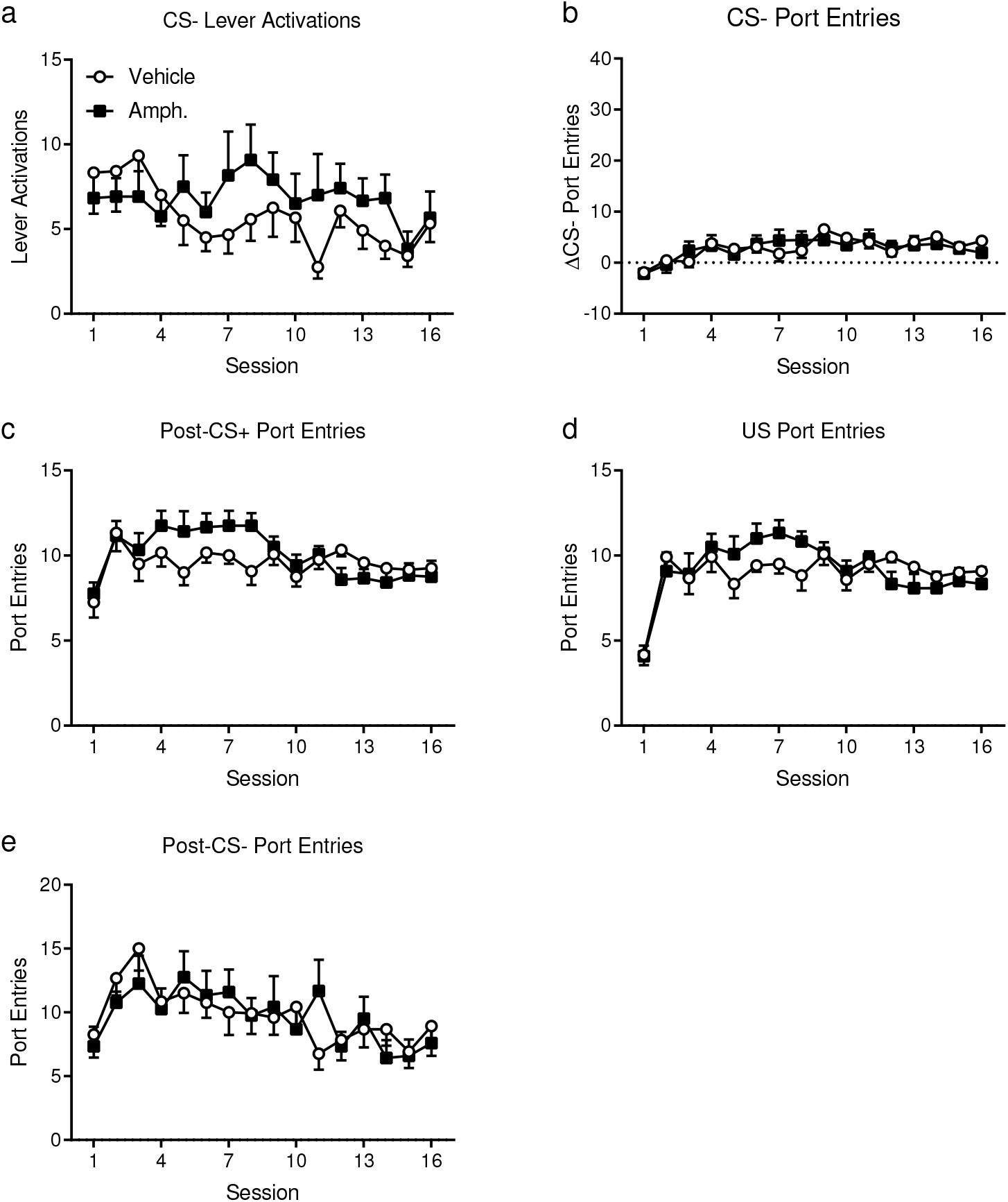
Responding elicited by the CS− and the Post-CS period during acquisition of sign-tracking. There were no differences between vehicle-treated and amphetamine-treated rats on (a) CS− lever activations, (b) ΔCS− port entries, (c) Post-CS+ port entries, (d) US port entries (i.e. the first 6 s of the Post-CS+ period during syringe pump operation) and (e) Post-CS− port entries. Data are means ± SEM. See Table A.6 for statistical results.

**Figure A.8.**
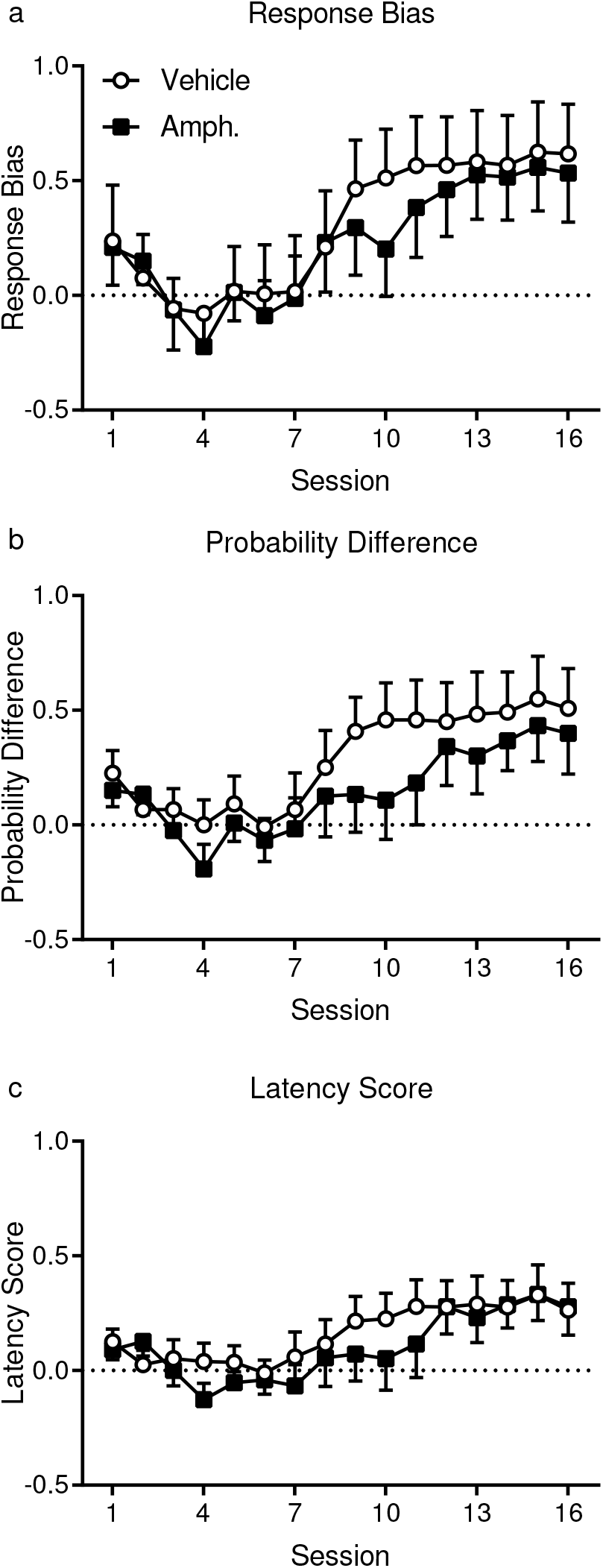
Component measures in the Pavlovian conditioned approach score. Prior amphetamine exposure did not alter (a) response bias, (b) probability difference or (c) latency score. Data are means ± SEM. See Table A.6 for statistical results.

**Table A.1.**
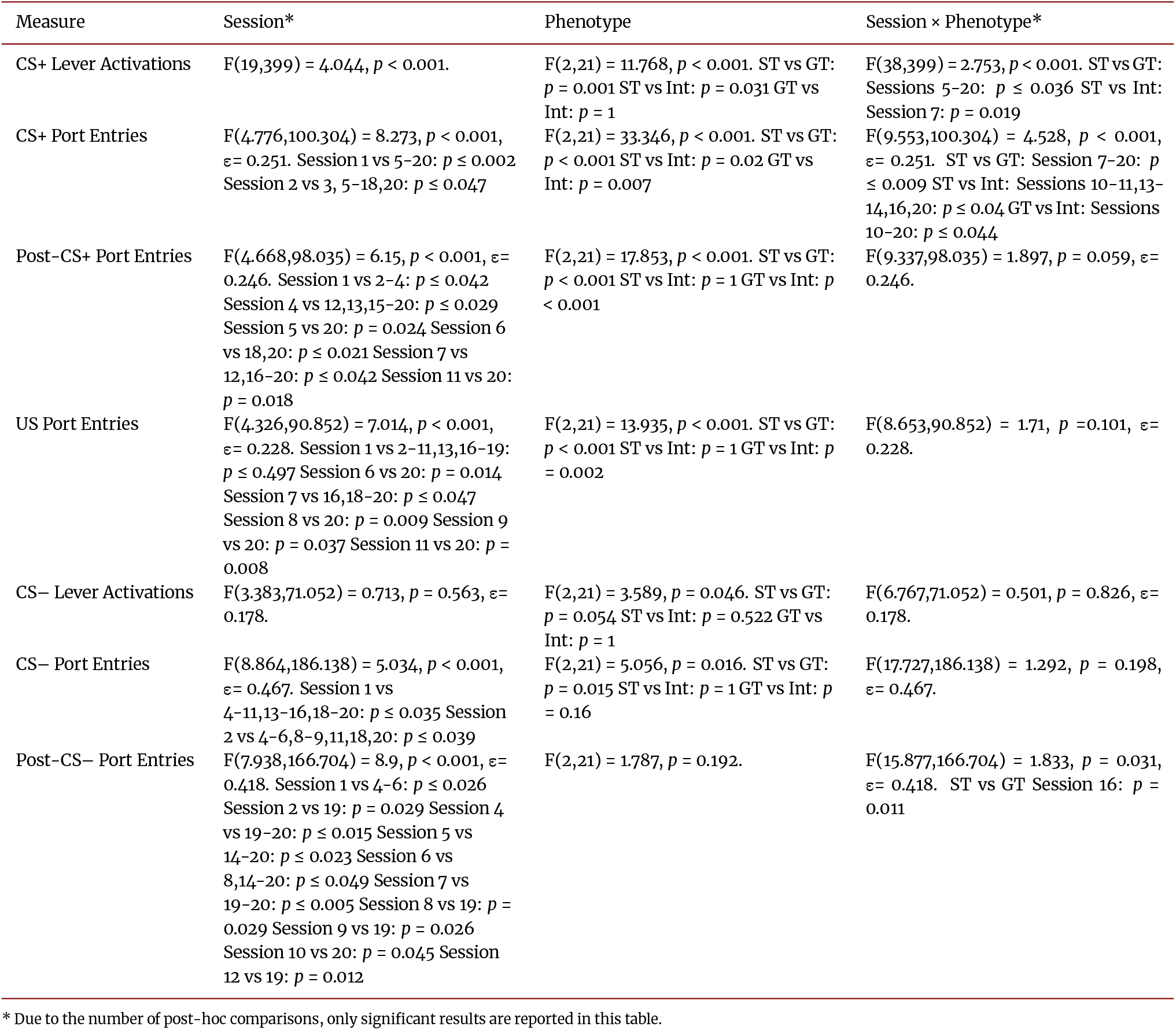
Statistical results for supplementary acquisition data.

**Table A.2.**
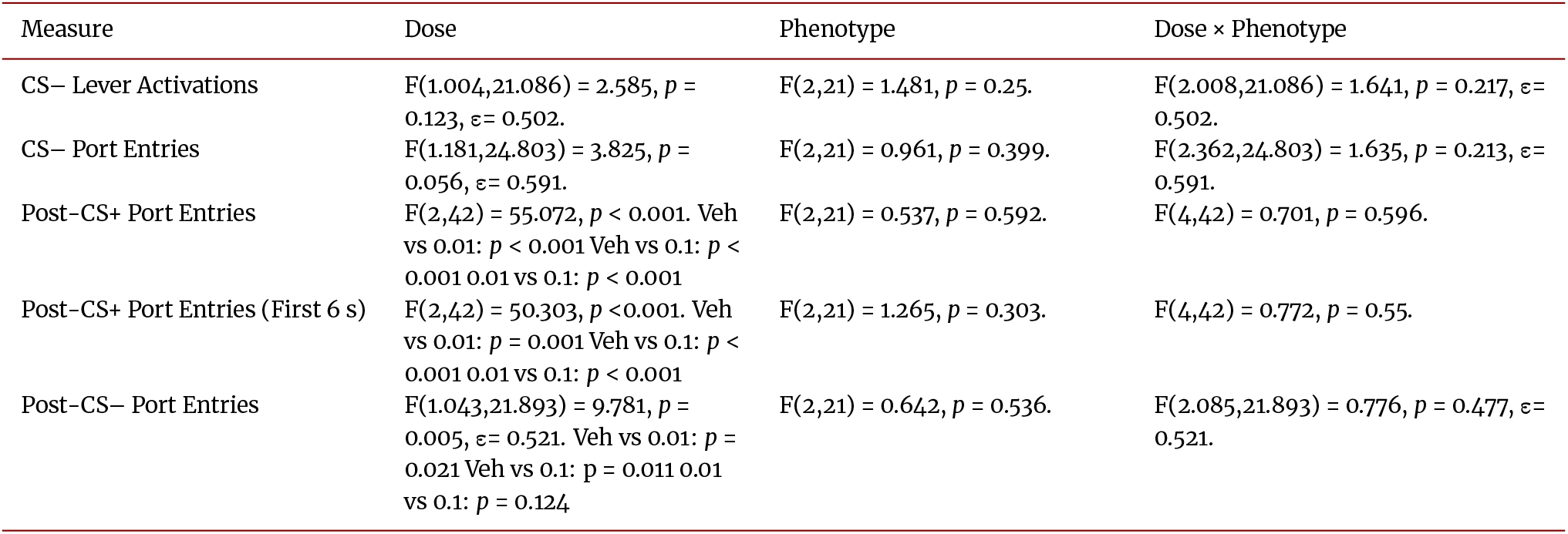
Statistical results for CS− and Post-CS measures during SCH-23390 tests.

**Table A.3.** Statistical results for supplementary measures during SCH-23390 tests.

| Measure | Dose | Phenotype | Dose × Phenotype |
| --- | --- | --- | --- |
| Probability CS+ Lever Activation | $F(1.377, 28.914) = 26.804, p < 0.001, \epsilon = 0.688$. Veh vs 0.01: $p = 0.002$; Veh vs 0.1: $p < 0.001$; 0.01 vs 0.1: $p = 0.001$. | $F(2, 21) = 7.984, p = 0.003$. ST vs GT: $p = 0.002$; ST vs Int: $p = 0.845$; GT vs Int: $p = 0.159$. | $F(2.754, 28.914) = 2.68, p = 0.07, \epsilon = 0.688$. |
| Latency CS+ Lever Activation | $F(1.199, 25.173) = 20.904, p < 0.001, \epsilon = 0.599$. Veh vs 0.01: $p = 0.002$; Veh vs 0.1: $p < 0.001$; 0.01 vs 0.1: $p = 0.004$. | $F(2, 21) = 4.374, p = 0.026$. ST vs GT: $p = 0.023$; ST vs Int: $p = 1$; GT vs Int: $p = 0.254$. | $F(2.397, 25.173) = 2.359, p = 0.107, \epsilon = 0.599$. |
| Probability CS+ Port Entry | $F(2, 42) = 26.186, p < 0.001$. Veh vs 0.01: $p = 0.002$; Veh vs 0.1: $p < 0.001$; 0.01 vs 0.1: $p = 0.004$. | $F(2, 21) = 5.325, p = 0.013$. ST vs GT: $p = 0.023$; ST vs Int: $p = 1$; GT vs Int: $p = 0.254$. | $F(4, 42) = 3.342, p = 0.018$. ST: Veh vs 0.01: $p = 0.42$; Veh vs 0.1: $p = 0.042$; 0.01 vs 0.1: $p = 0.405$. GT: Veh vs 0.01: $p = 0.034$; Veh vs 0.1: $p < 0.001$; 0.01 vs 0.1: $p = 0.003$. Int: Veh vs 0.01: $p = 0.67$; Veh vs 0.1: $p = 0.009$; 0.01 vs 0.1: $p = 0.025$. |
| Latency CS+ Port Entry | $F(2, 42) = 18.005, p < 0.001$. Veh vs 0.01: $p = 0.26$; Veh vs 0.1: $p < 0.001$; 0.01 vs 0.1: $p < 0.001$. | $F(2, 21) = 7.441, p = 0.004$. ST vs GT: $p = 0.26$; ST vs Int: $p < 0.001$; GT vs Int: $p < 0.001$. | $F(4, 42) = 3.058, p = 0.027$. ST: Veh vs 0.01: $p = 0.865$; Veh vs 0.1: $p = 0.088$; 0.01 vs 0.1: $p = 0.518$. GT: Veh vs 0.01: $p = 0.311$; Veh vs 0.1: $p < 0.001$; 0.01 vs 0.1: $p < 0.001$. Int: Veh vs 0.01: $p = 1$; Veh vs 0.1: $p = 0.075$; 0.01 vs 0.1: $p = 0.089$. |
| PCA Score | $F(1.338, 28.095) = 0.625, p = 0.481, \epsilon = 0.669$. | $F(2, 21) = 14.103, p < 0.001$. ST vs GT: $p < 0.001$; ST vs Int: $p = 0.249$; GT vs Int: $p = 0.077$. | $F(2.676, 28.095) = 5.381, p = 0.006$. ST: Veh vs 0.01: $p = 0.753$; Veh vs 0.1: $p = 0.004$; 0.01 vs 0.1: $p = 0.001$. GT: Veh vs 0.01: $p = 1$; Veh vs 0.1: $p = 0.065$; 0.01 vs 0.1: $p = 0.001$. Int: Veh vs 0.01: $p = 1$; Veh vs 0.1: $p = 1$; 0.01 vs 0.1: $p = 1$. |
| Response Bias | $F(2, 42) = 14.484, p < 0.001$. Veh vs 0.01: $p = 1$; Veh vs 0.1: $p < 0.001$; 0.01 vs 0.1: $p < 0.001$. | $F(2, 21) = 12.135, p < 0.001$. ST vs GT: $p < 0.001$; ST vs Int: $p = 0.191$; GT vs Int: $p = 0.189$. | $F(4, 42) = 2.606, p = 0.049$. ST: Veh vs 0.01: $p = 1$; Veh vs 0.1: $p < 0.001$; 0.01 vs 0.1: $p < 0.001$. GT: Veh vs 0.01: $p = 0.81$; Veh vs 0.1: $p = 0.423$; 0.01 vs 0.1: $p = 1$. Int: Veh vs 0.01: $p = 1$; Veh vs 0.1: $p = 0.082$; 0.01 vs 0.1: $p = 0.014$. |
| Probability Difference | $F(1.371, 28.781) = 0.596, p = 0.497, \epsilon = 0.685$. | $F(2, 21) = 13.793, p < 0.001$. ST vs GT: $p < 0.001$; ST vs Int: $p = 0.123$; GT vs Int: $p = 0.174$. | $F(2.741, 28.781) = 5.617, p = 0.005, \epsilon = 0.685$. ST: Veh vs 0.01: $p = 0.148$; Veh vs 0.1: $p = 0.004$; 0.01 vs 0.1: $p = 0.042$. GT: Veh vs 0.01: $p = 1$; Veh vs 0.1: $p = 0.042$; 0.01 vs 0.1: $p = 0.01$. Int: Veh vs 0.01: $p = 1$; Veh vs 0.1: $p = 1$; 0.01 vs 0.1: $p = 1$. |
| Latency Score | $F(1.321, 27.749) = 1.674, p = 0.209, \epsilon = 0.661$. | $F(2, 21) = 7.441, p = 0.004$. ST vs GT: $p < 0.001$; ST vs Int: $p = 0.689$; GT vs Int: $p = 0.071$. | $F(2.643, 27.749) = 4.174, p = 0.018, \epsilon = 0.661$. ST: Veh vs 0.01: $p = 0.105$; Veh vs 0.1: $p = 0.02$; 0.01 vs 0.1: $p = 0.754$. GT: Veh vs 0.01: $p = 1$; Veh vs 0.1: $p = 0.082$; 0.01 vs 0.1: $p = 0.002$. Int: Veh vs 0.01: $p = 0.786$; Veh vs 0.1: $p = 1$; 0.01 vs 0.1: $p = 1$. |

**Table A.4.** Statistical results for CS− and Post-CS measures during eticlopride tests.

| Measure | Dose | Phenotype | Dose × Phenotype |
| --- | --- | --- | --- |
| CS– Lever Activations | $F(1.245, 26.145) = 3.914, p = 0.0504, \epsilon = 0.623.$ | $F(2, 21) = 0.638, p = 0.538.$ | $F(2.49, 26.145) = 0.3, p = 0.789, \epsilon = 0.623.$ |
| CS– Port Entries | $F(2, 42) = 1.722, p = 0.191.$ | $F(2, 21) = 1.098, p = 0.352.$ | $F(4, 42) = 0.79, p = 0.538.$ |
| Post-CS+ Port Entries | $F(2, 42) = 132.57, p < 0.001.$ Veh vs 0.01: $p = 0.229$ Veh vs 0.1: $p < 0.001$ 0.01 vs 0.1: $p < 0.001$ | $F(2, 21) = 0.291, p = 0.751.$ | $F(4, 42) = 0.835, p = 0.51.$ |
| Post-CS+ Port Entries (First 6 s) | $F(2, 42) = 136.693, p < 0.001.$ Veh vs 0.01: $p = 0.353$ Veh vs 0.1: $p < 0.001$ 0.01 vs 0.1: $p < 0.001$ | $F(2, 21) = 0.467, p = 0.633.$ | $F(4, 42) = 0.35, p = 0.842.$ |
| Post-CS– Port Entries | $F(2, 42) = 8.431, p = 0.001.$ Veh vs 0.01: $p = 1$ Veh vs 0.1: $p = 0.001$ 0.01 vs 0.1: $p = 0.007$ | $F(2, 21) = 1.308, p = 0.292.$ | $F(4, 42) = 0.751, p = 0.563.$ |

**Table A.5.**
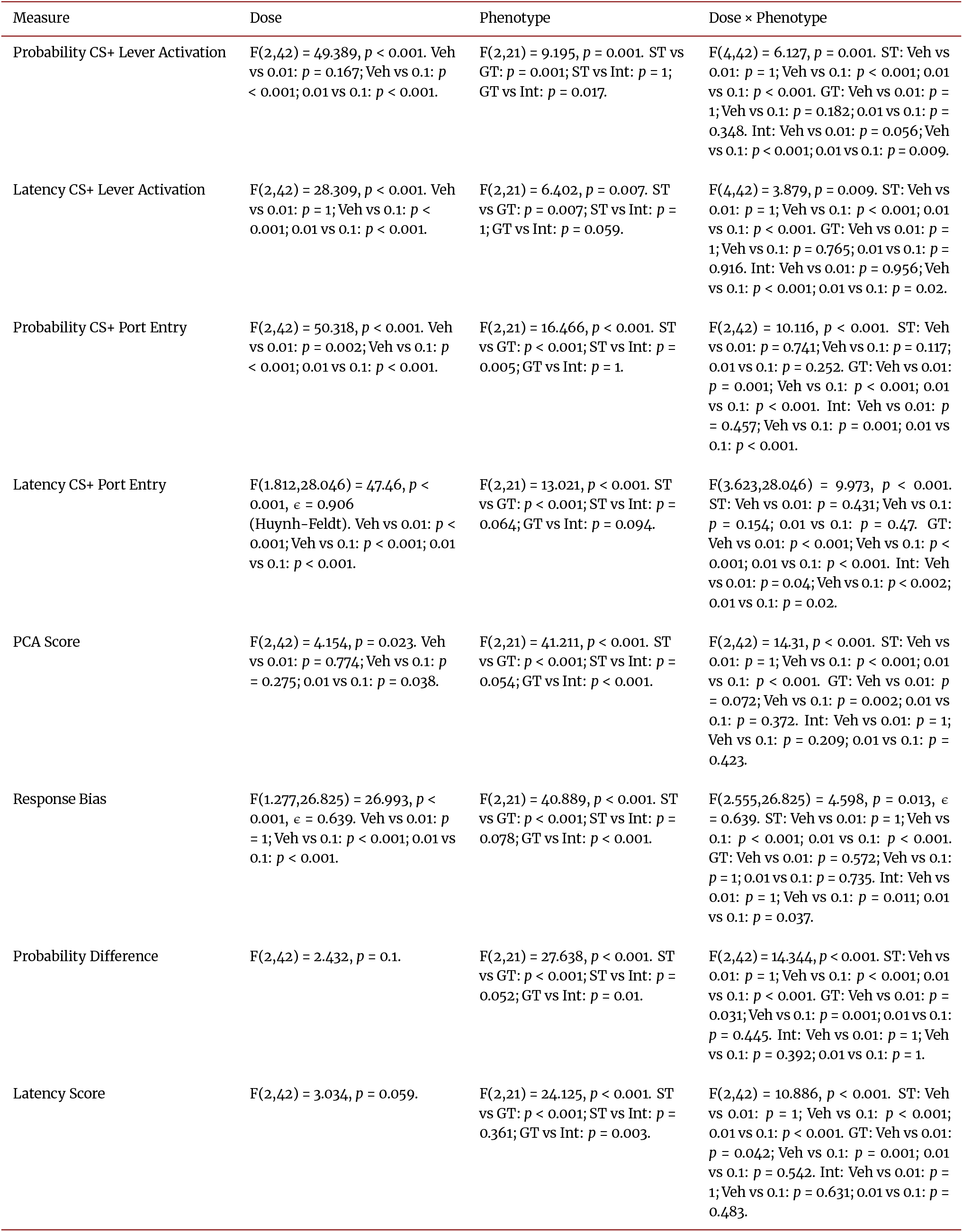
Statistical results for supplementary measures from eticlopride tests.

| Measure | Dose | Phenotype | Dose × Phenotype |
| --- | --- | --- | --- |
| Probability CS+ Lever Activation | $F(2,42) = 49.389, p < 0.001$. Veh vs 0.01: $p = 0.167$; Veh vs 0.1: $p < 0.001$; 0.01 vs 0.1: $p < 0.001$. | $F(2,21) = 9.195, p = 0.001$. ST vs GT: $p = 0.001$; ST vs Int: $p = 1$; GT vs Int: $p = 0.017$. | $F(4,42) = 6.127, p = 0.001$. ST: Veh vs 0.01: $p = 1$; Veh vs 0.1: $p < 0.001$; 0.01 vs 0.1: $p < 0.001$. GT: Veh vs 0.01: $p = 1$; Veh vs 0.1: $p = 0.182$; 0.01 vs 0.1: $p = 0.348$. Int: Veh vs 0.01: $p = 0.056$; Veh vs 0.1: $p < 0.001$; 0.01 vs 0.1: $p = 0.009$. |
| Latency CS+ Lever Activation | $F(2,42) = 28.309, p < 0.001$. Veh vs 0.01: $p = 1$; Veh vs 0.1: $p < 0.001$; 0.01 vs 0.1: $p < 0.001$. | $F(2,21) = 6.402, p = 0.007$. ST vs GT: $p = 0.007$; ST vs Int: $p = 1$; GT vs Int: $p = 0.059$. | $F(4,42) = 3.879, p = 0.009$. ST: Veh vs 0.01: $p = 1$; Veh vs 0.1: $p < 0.001$; 0.01 vs 0.1: $p < 0.001$. GT: Veh vs 0.01: $p = 1$; Veh vs 0.1: $p = 0.765$; 0.01 vs 0.1: $p = 0.916$. Int: Veh vs 0.01: $p = 0.956$; Veh vs 0.1: $p < 0.001$; 0.01 vs 0.1: $p = 0.02$. |
| Probability CS+ Port Entry | $F(2,42) = 50.318, p < 0.001$. Veh vs 0.01: $p = 0.002$; Veh vs 0.1: $p < 0.001$; 0.01 vs 0.1: $p < 0.001$. | $F(2,21) = 16.466, p < 0.001$. ST vs GT: $p < 0.001$; ST vs Int: $p = 0.005$; GT vs Int: $p = 1$. | $F(2,42) = 10.116, p < 0.001$. ST: Veh vs 0.01: $p = 0.741$; Veh vs 0.1: $p = 0.117$; 0.01 vs 0.1: $p = 0.252$. GT: Veh vs 0.01: $p = 0.001$; Veh vs 0.1: $p < 0.001$; 0.01 vs 0.1: $p < 0.001$. Int: Veh vs 0.01: $p = 0.457$; Veh vs 0.1: $p = 0.001$; 0.01 vs 0.1: $p < 0.001$. |
| Latency CS+ Port Entry | $F(1.812, 28.046) = 47.46, p < 0.001, \epsilon = 0.906$ (Huynh–Feldt). Veh vs 0.01: $p < 0.001$; Veh vs 0.1: $p < 0.001$; 0.01 vs 0.1: $p < 0.001$. | $F(2,21) = 13.021, p < 0.001$. ST vs GT: $p < 0.001$; ST vs Int: $p = 0.064$; GT vs Int: $p = 0.094$. | $F(3.623, 28.046) = 9.973, p < 0.001$. ST: Veh vs 0.01: $p = 0.431$; Veh vs 0.1: $p = 0.154$; 0.01 vs 0.1: $p = 0.47$. GT: Veh vs 0.01: $p < 0.001$; Veh vs 0.1: $p < 0.001$; 0.01 vs 0.1: $p < 0.001$. Int: Veh vs 0.01: $p = 0.04$; Veh vs 0.1: $p < 0.002$; 0.01 vs 0.1: $p = 0.02$. |
| PCA Score | $F(2,42) = 4.154, p = 0.023$. Veh vs 0.01: $p = 0.774$; Veh vs 0.1: $p = 0.275$; 0.01 vs 0.1: $p = 0.038$. | $F(2,21) = 41.211, p < 0.001$. ST vs GT: $p < 0.001$; ST vs Int: $p = 0.054$; GT vs Int: $p < 0.001$. | $F(2,42) = 14.31, p < 0.001$. ST: Veh vs 0.01: $p = 1$; Veh vs 0.1: $p < 0.001$; 0.01 vs 0.1: $p < 0.001$. GT: Veh vs 0.01: $p = 0.072$; Veh vs 0.1: $p = 0.002$; 0.01 vs 0.1: $p = 0.372$. Int: Veh vs 0.01: $p = 1$; Veh vs 0.1: $p = 0.209$; 0.01 vs 0.1: $p = 0.423$. |
| Response Bias | $F(1.277, 26.825) = 26.993, p < 0.001, \epsilon = 0.639$. Veh vs 0.01: $p = 1$; Veh vs 0.1: $p < 0.001$; 0.01 vs 0.1: $p < 0.001$. | $F(2,21) = 40.889, p < 0.001$. ST vs GT: $p < 0.001$; ST vs Int: $p = 0.078$; GT vs Int: $p < 0.001$. | $F(2.555, 26.825) = 4.598, p = 0.013, \epsilon = 0.639$. ST: Veh vs 0.01: $p = 1$; Veh vs 0.1: $p < 0.001$; 0.01 vs 0.1: $p < 0.001$. GT: Veh vs 0.01: $p = 0.572$; Veh vs 0.1: $p = 1$; 0.01 vs 0.1: $p = 0.735$. Int: Veh vs 0.01: $p = 1$; Veh vs 0.1: $p = 0.011$; 0.01 vs 0.1: $p = 0.037$. |
| Probability Difference | $F(2,42) = 2.432, p = 0.1$. | $F(2,21) = 27.638, p < 0.001$. ST vs GT: $p < 0.001$; ST vs Int: $p = 0.052$; GT vs Int: $p = 0.01$. | $F(2,42) = 14.344, p < 0.001$. ST: Veh vs 0.01: $p = 1$; Veh vs 0.1: $p < 0.001$; 0.01 vs 0.1: $p < 0.001$. GT: Veh vs 0.01: $p = 0.031$; Veh vs 0.1: $p = 0.001$; 0.01 vs 0.1: $p = 0.445$. Int: Veh vs 0.01: $p = 1$; Veh vs 0.1: $p = 0.392$; 0.01 vs 0.1: $p = 1$. |
| Latency Score | $F(2,42) = 3.034, p = 0.059$. | $F(2,21) = 24.125, p < 0.001$. ST vs GT: $p < 0.001$; ST vs Int: $p = 0.361$; GT vs Int: $p = 0.003$. | $F(2,42) = 10.886, p < 0.001$. ST: Veh vs 0.01: $p = 1$; Veh vs 0.1: $p < 0.001$; 0.01 vs 0.1: $p < 0.001$. GT: Veh vs 0.01: $p = 0.042$; Veh vs 0.1: $p = 0.001$; 0.01 vs 0.1: $p = 0.542$. Int: Veh vs 0.01: $p = 1$; Veh vs 0.1: $p = 0.631$; 0.01 vs 0.1: $p = 0.483$. |

**Table A.6.**
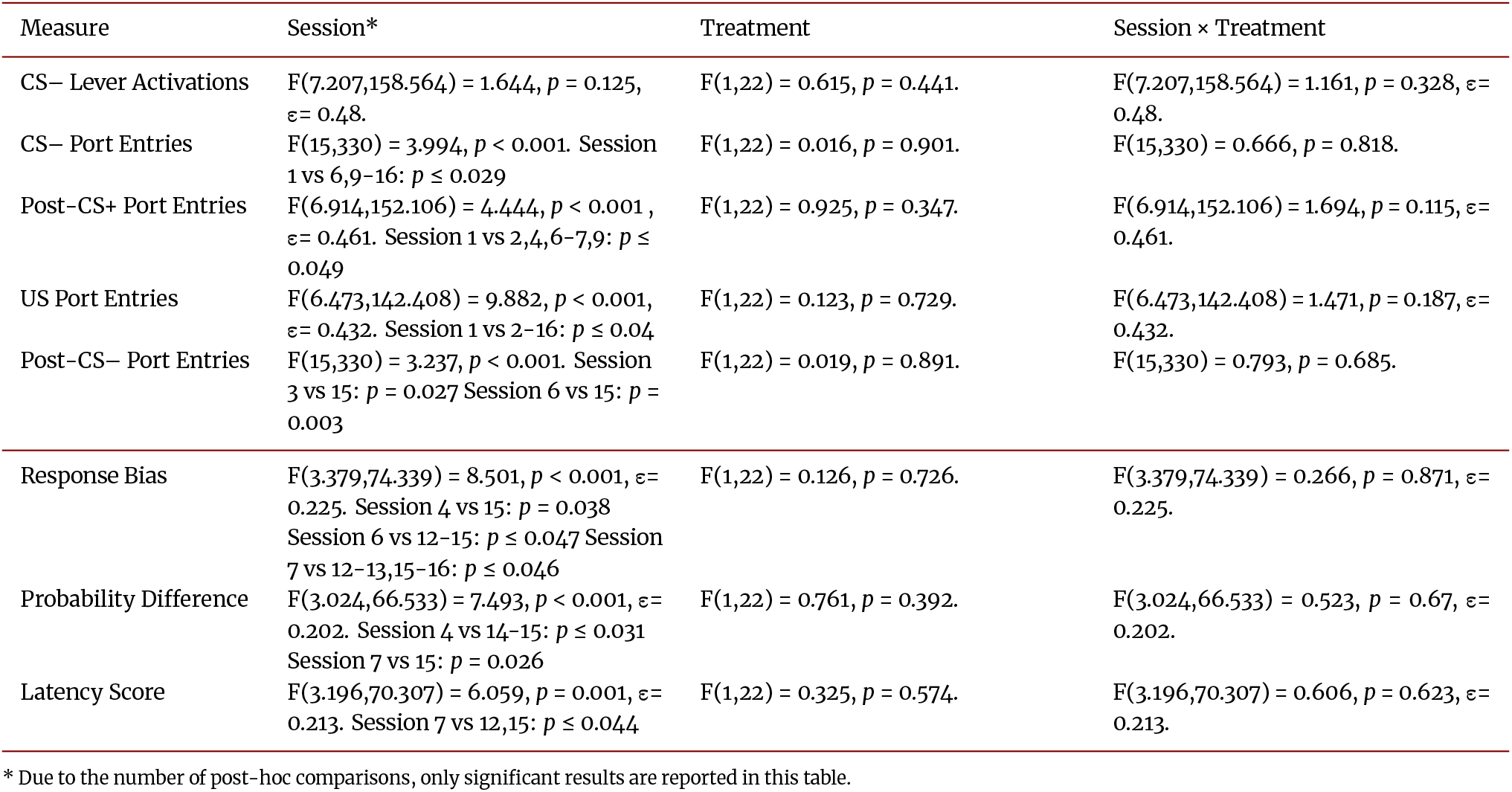
Statistical results for supplementary measures and PCA score components during acquisition of sign-tracking.

| Measure | Session* | Treatment | Session × Treatment |
| --- | --- | --- | --- |
| CS– Lever Activations | $F(7.207,158.564) = 1.644, p = 0.125, \epsilon = 0.48.$ | $F(1,22) = 0.615, p = 0.441.$ | $F(7.207,158.564) = 1.161, p = 0.328, \epsilon = 0.48.$ |
| CS– Port Entries | $F(15,330) = 3.994, p < 0.001.$ Session 1 vs 6,9–16: $p \leq 0.029$ | $F(1,22) = 0.016, p = 0.901.$ | $F(15,330) = 0.666, p = 0.818.$ |
| Post-CS+ Port Entries | $F(6.914,152.106) = 4.444, p < 0.001, \epsilon = 0.461.$ Session 1 vs 2,4,6–7,9: $p \leq 0.049$ | $F(1,22) = 0.925, p = 0.347.$ | $F(6.914,152.106) = 1.694, p = 0.115, \epsilon = 0.461.$ |
| US Port Entries | $F(6.473,142.408) = 9.882, p < 0.001, \epsilon = 0.432.$ Session 1 vs 2–16: $p \leq 0.04$ | $F(1,22) = 0.123, p = 0.729.$ | $F(6.473,142.408) = 1.471, p = 0.187, \epsilon = 0.432.$ |
| Post-CS– Port Entries | $F(15,330) = 3.237, p < 0.001.$ Session 3 vs 15: $p = 0.027$ Session 6 vs 15: $p = 0.003$ | $F(1,22) = 0.019, p = 0.891.$ | $F(15,330) = 0.793, p = 0.685.$ |
| Response Bias | $F(3.379,74.339) = 8.501, p < 0.001, \epsilon = 0.225.$ Session 4 vs 15: $p = 0.038$ Session 6 vs 12–15: $p \leq 0.047$ Session 7 vs 12–13,15–16: $p \leq 0.046$ | $F(1,22) = 0.126, p = 0.726.$ | $F(3.379,74.339) = 0.266, p = 0.871, \epsilon = 0.225.$ |
| Probability Difference | $F(3.024,66.533) = 7.493, p < 0.001, \epsilon = 0.202.$ Session 4 vs 14–15: $p \leq 0.031$ Session 7 vs 15: $p = 0.026$ | $F(1,22) = 0.761, p = 0.392.$ | $F(3.024,66.533) = 0.523, p = 0.67, \epsilon = 0.202.$ |
| Latency Score | $F(3.196,70.307) = 6.059, p = 0.001, \epsilon = 0.213.$ Session 7 vs 12,15: $p \leq 0.044$ | $F(1,22) = 0.325, p = 0.574.$ | $F(3.196,70.307) = 0.606, p = 0.623, \epsilon = 0.213.$ |

\* Due to the number of post-hoc comparisons, only significant results are reported in this table.

